# Detecting changes in dynamical structures in synchronous neural oscillations using probabilistic inference

**DOI:** 10.1101/2020.10.13.335356

**Authors:** Hiroshi Yokoyama, Keiichi Kitajo

**Affiliations:** Division of Neural Dynamics, Department of System Neuroscience, National Institute for Physiological Sciences, National Institutes of Natural Sciences, Okazaki 444-8585, Japan; Department of Physiological Sciences, School of Life Science, Graduate University for Advanced Studies (SOKENDAI), Okazaki 444-8585, Japan; CBS-TOYOTA Collaboration Center, RIKEN Center for Brain Science, Wako, 351-0198, Japan

**Keywords:** Bayesian inference, change point detection, Kullback-Leibler divergence, phase-coupled oscillator model, electroencephalography

## Abstract

Recent neuroscience studies have suggested that cognitive functions and learning capacity are reflected in the time-evolving dynamics of brain networks. However, an efficient method to detect changes in dynamical brain structures using neural data has yet to be established. To address this issue, we developed a new model-based approach to detect change points in dynamical network structures by combining the model-based network estimation with a phase-coupled oscillator model and sequential Bayesian inference. By giving the model parameter as the prior distribution, applying Bayesian inference allows the extent of temporal changes in dynamic brain networks to be quantified by comparing the prior distribution with posterior distribution using information theoretical criteria. For this, we used the Kullback-Leibler divergence as an index of such changes. To validate our method, we applied it to numerical data and electroencephalography data. As a result, we confirmed that the Kullback-Leibler divergence only increased when changes in dynamical network structures occurred. Our proposed method successfully estimated both directed network couplings and change points of dynamical structures in the numerical and electroencephalography data. These results suggest that our proposed method can reveal the neural basis of dynamic brain networks.

## 1. Introduction

The ability to detect temporal changes of network dynamics from neural data has become of utmost importance for elucidating the role of neural dynamics in the brain. For instance, many neuroscience studies on the functional role of network dynamics in the brain have discussed the relevance of nonlinear coupled dynamical systems (Penny et al., 2009; Breakspear et al., 2010; Cabral et al., 2011; Deco et al., 2017b,a; Sase & Kitajo, 2021). In addition, some recent experimental studies have raised the possibility that flexible changes in dynamic brain networks could be associated with cognitive functions and learning capacity (Bassett et al., 2011; Baum et al., 2017; Braun et al., 2015, 2016). However, an efficient method to detect changes in dynamical brain structures using neural data has yet to be established.

To detect temporal changes in brain networks in a data-driven manner, both temporal dependence in time-series data and structural relationships within the whole brain should be considered. In general, multivariate time-series data with non-stationary dynamics, such as neural data, not only contain information about structural relationships between dataset, but also about temporal dependence (i.e., the causal relationship from past to current states). Most previous studies have evaluated time-varying functional connectivity by applying the sliding window method (Hutchison et al., 2013; Damaraju et al., 2014). In this method, time series data are transformed into short epochs, each of which is analyzed separately and without explicitly dealing with the time-evolving dynamics. In most studies that have applied functional connectivity analysis, when evaluating the network structure within each time window, the coupling strength for each pair of brain areas is calculated based on the pairwise correlation between the time-series data. According to prior evidence from an economic study (Granger et al., 1974), correlation coefficients between two different time-series data with a high auto-correlation tend to be large, regardless of the existence of actual causality between the two time-series data. Therefore, the conventional method of functional connectivity analysis cannot efficiently quantify the changes in time-evolving network dynamics in the brain. To tackle this issue, methodological studies have proposed data-driven approaches to measure time-varying functional connectivity based on the adaptive auto-regressive model (Astolfi et al., 2008; Leistritz et al., 2013). However, these models are not sufficient considering the nonlinear dynamical properties of brain networks, because they assume that both temporal dynamics and network interactions are linear. One possible approach to avoid the above-mentioned issues is to utilize a dynamical model-based network analysis. Namely, applying a coupled dynamical model for brain network analysis would allow the consideration of both network couplings and time-evolving dynamics reflected in neuronal time-series data. Given that neural data contain a strong temporal dependence in time-evolving dynamics, as well as network coupling, making the assumption that a dynamical system underlies neural data could be an essential way to address the aforementioned issues associated with functional connectivity analyses. As an additional advantage, this method would also reduce spurious network couplings, because a model-based analysis can differentiate between direct and indirect couplings.

However, when using a model-based approach, it is important to select a model that can estimate network couplings, because the model must be able to accurately explain the dynamical properties of the observed time-series data. To address the above issues, we developed a novel model-based method to detect temporal changes in dynamic brain networks that combines sequential Bayesian inference and dynamical model-based network analysis. Based on studies on nonlinear brain dynamics (Penny et al., 2009; Breakspear et al., 2010; Cabral et al., 2011; Deco et al., 2017b,a), we assumed that the time-evolving dynamics and the data structures of brain networks could be expressed as a phase-coupled oscillator model in this proposed method (Netoff et al., 2012; Penny et al., 2009; Ota & Aoyagi, 2014; Suzuki et al., 2018; Onojima et al., 2018). Moreover, to capture the time-varying changes in network dynamics, our method recursively fitted the observed time series data to the phase-coupled oscillator model using sequential Bayesian inference (Sarris, 1973; Bishop, 2007). The advantage of using sequential Bayesian inference for a model fitting is that, by giving the model parameter as the prior distribution, the extent of temporal changes in the brain network can be quantified based on the comparison between prior and posterior distributions using information theoretical criteria. Considering this, we predicted that temporal changes in dynamical structures of the brain network would be reflected in the change in model statistics regarding the prior distribution. Therefore, to define an index for the change in dynamical structures of functional brain networks, the difference between prior and posterior distributions of model parameters was measured using the Kullback-Leibler (KL) divergence.

To validate our proposed method, two examinations were performed. First, we applied our method to a time series of synthetic data with known dynamical changes of network coupling, which were generated by a weakly phase-coupled oscillator model. Doing so allowed us to confirm whether the change points detected by our proposed method were consistent with the exact timing of network coupling changes in the synthetic data. Next, we applied our method to an open dataset of real electroencephalography (EEG) signals measured in healthy human participants who were passively listening to binaural auditory stimuli of 40-Hz amplitude-modulated (AM) white noise (McFadden et al., 2014). In this way, we assessed whether our proposed method could detect the timing of changes in phase-coupled EEG networks induced by the periodic auditory stimuli.

## 2. Materials and methods

In this section, we will first provide an overview of our proposed method and the details of its algorithm. We will then explain the procedures for model validation using synthetic data from numerical models and EEG data.

### 2.1. An overview of the proposed method

In our proposed method, to estimate the dynamical structure in the brain networks, we applied a data-driven modeling with a phase-coupled oscillator (Kuramoto, 1984; Ko & Ermentrout, 2009; Netoff et al., 2012; Stankovski et al., 2017; Pietras & Daffertshofer, 2019; Onojima et al., 2018; Suzuki et al., 2018; Ota & Aoyagi, 2014). From the point of view of the dynamical systems theory, if a dynamical system exhibits limit cycles with a stable point, the system can be theoretically described by a phase *ϕ* as a simple dynamical system with one degree of freedom (Kuramoto, 1984; Pietras & Daffertshofer, 2019; Onojima et al., 2018; Suzuki et al., 2018; Ota & Aoyagi, 2014). Furthermore, when dynamical systems weakly interact with each other around the stable limit cycle attractors, this system can be described as a phase-coupled oscillator model. We have provided a more detailed description of the background of this theory in the Supplementary Materials. Considering this, we presume that phase-coupled oscillator dynamics underlie synchronous networks in the brain. Here, we assume that the time derivative of phase oscillator 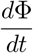 can be described with the function *F* (**Φ**, *t*; ***K***) characterized by the phase **Φ** at time *t* and the model parameter ***K*** (i.e., 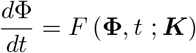). Furthermore, if the model parameter ***K*** is associated with network couplings, the temporal changes in brain network dynamics will be mediated by the changes of the model parameters ***K***. Based on this assumption, we proposed the following new method for the network change point detection, as shown in Figure 1. In our proposed method, to capture the temporal changes in the model parameter ***K***, the phase of the measured data is fitted with a phase-coupled oscillator model on a sample-by-sample basis. Given the prior distribution of the model parameters, the parameter ***K*** can be calculated using the Bayesian rule, as shown in Figure 1. As the advantage of using sequential Bayesian inference for a model fitting, by giving the model parameter as the prior distribution, the extent of temporal changes in the brain network can be quantified based on the comparison between prior and posterior distributions using information theoretical criteria. Therefore, the changes in dynamical structures of brain networks are quantified using the Kullback-Leibler divergence between prior and posterior distributions on a sample-by-sample basis (Figure 1).

**Figure 1:**
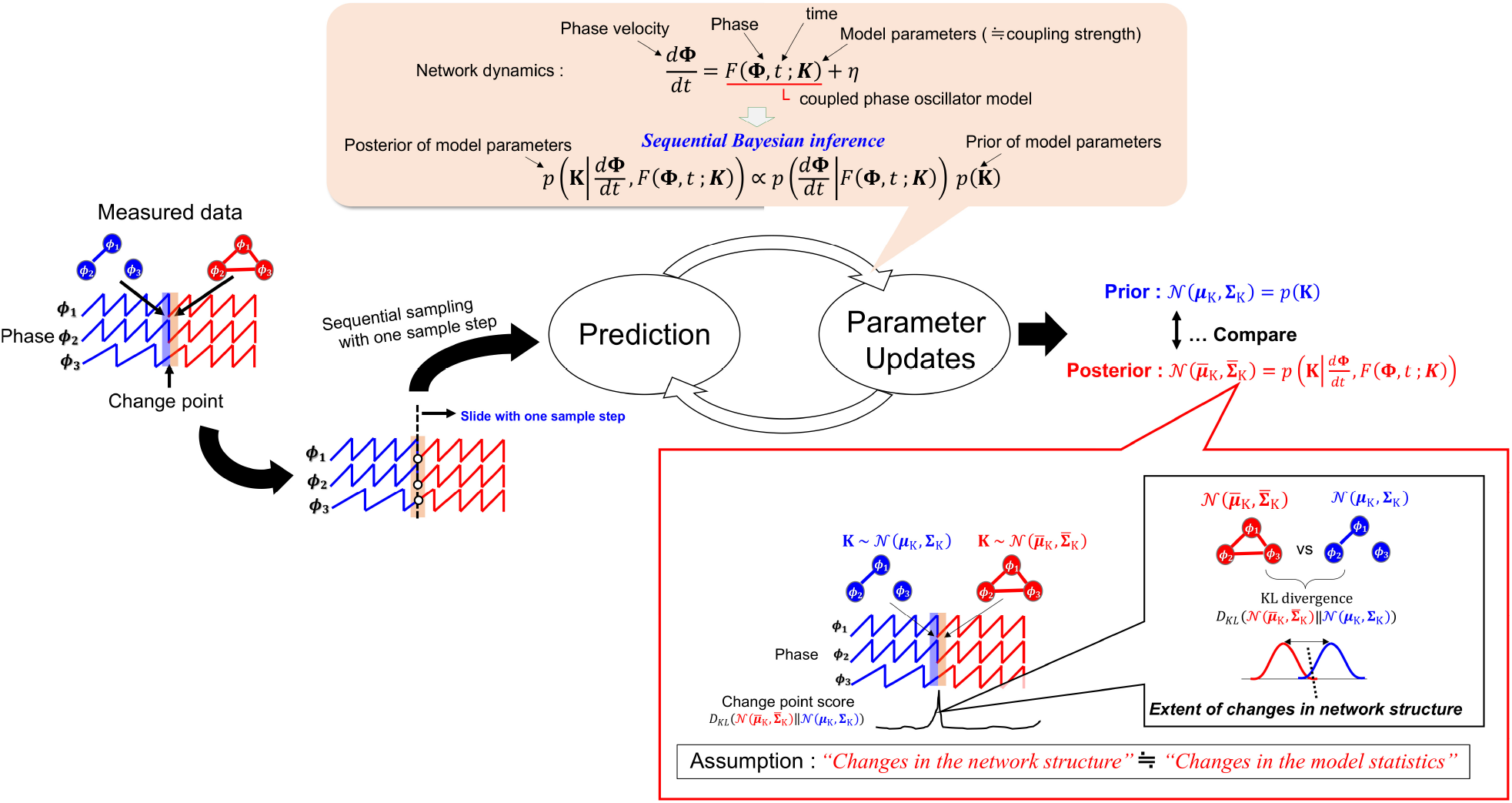
Overview of the proposed method for network change point detection. In the proposed method, the time series data are fitted with a phase-coupled oscillator model using Bayesian inference. Here we assume that the changes in dynamical networks are mediated by the changes of the model parameters regarding the coupling strengths between phase oscillators. By giving the prior distribution of the model parameters, the structural changes in networks can be quantified by comparing prior and updated posterior distributions. Therefore, the changes in dynamical structures of brain networks are quantified using the Kullback-Leibler divergence between prior and posterior distributions. In the proposed method, by using sequential Bayesian inference, the model parameter updates are conducted on a sample-by-sample basis, and the metric of change point score using the KL divergence is also calculated after updating the prior of the model parameters sample-by-sample.

### 2.2. Estimation of the network coupling and change point scores

In this section, we describe how the model parameter estimation and change point detection are conducted. As mentioned above, for our proposed method, we consider the phase-coupled oscillators as the underlying dynamical system in the brain networks. Based on the dynamical systems theory, an *N*-th phase-coupled oscillator model can be described using the following equation (see the Supplementary Materials for a detailed description of the theoretical background):

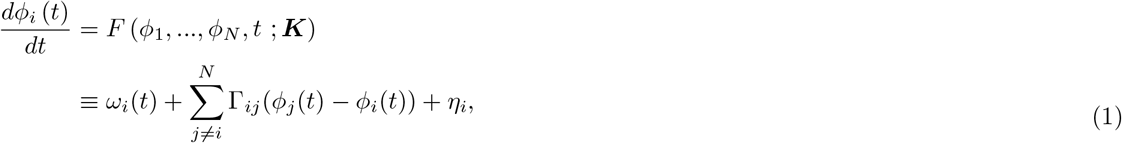

where,

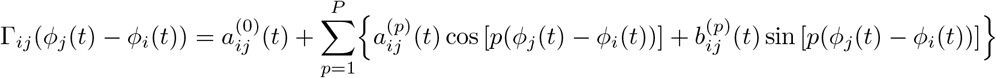

Note that *ϕ_i_*(*t*), *ω_i_*(*t*), and *η_i_*(*t*) indicate the phase, the natural frequency, and the observation noise of the *i*-th oscillator, respectively. Moreover, the function Γ_*ij*_ (*ϕ_j_* – *ϕ_i_*) reflects the extent of the coupling between oscillators *i*, *j*. In our study, as in some computational and methodological neuroscience studies (Penny et al., 2009; Netoff et al., 2012; Ota & Aoyagi, 2014; Suzuki et al., 2018; Onojima et al., 2018), given that the function Γ_*ij*_ (*ϕ_j_*(*t*) – *ϕ_i_*(*t*)) is a 2-*π* periodic function when there exists a limit-cycle attractor of the dynamical system, we approximated the function Γ_*ij*_ (*ϕ_j_*(*t*) – *ϕ_i_*(*t*)) to the *P*-th Fourier series, as follows:

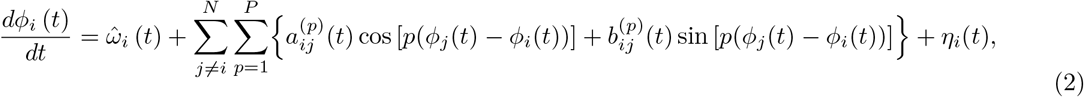

where

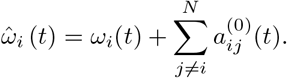

Note that 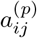 and 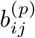 indicate the coefficients of the *p*-th Fourier series corresponding to the coupling strength between the *i* and *j*-th oscillators. Moreover, 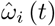 is the neutral frequency of this model. Therefore, in our method, the network dynamics in the brain can be considered as the function that is explained using the phase interaction Γ_*ij*_ (*ϕ_j_*(*t*) – *ϕ_i_*(*t*)). Moreover, the Γ_*ij*_ (*ϕ_j_* – *ϕ_i_*) is approximated by the *p*-th Fourier series parameterized by the coefficients 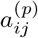 and 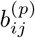. In our proposed method, the coupling strength *C_ij_* for all *i*, *j* is defined as follows:

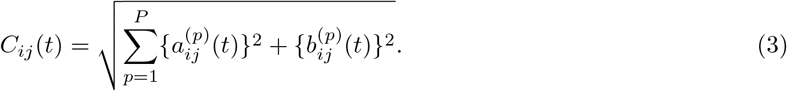

To evaluate the temporal changes in the networks, we extended the parameter estimation algorithm in Eq. (2) using Bayesian inference (Bishop, 2007; Sarris, 1973). In this way, the parameters of Eq. (2) could be directly estimated from the observed data in the same way as for sequential Bayesian regression. To apply Bayesian inference, Eq. (2) was rewritten in its matrix form, as shown in Eq. (4).

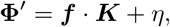

where,

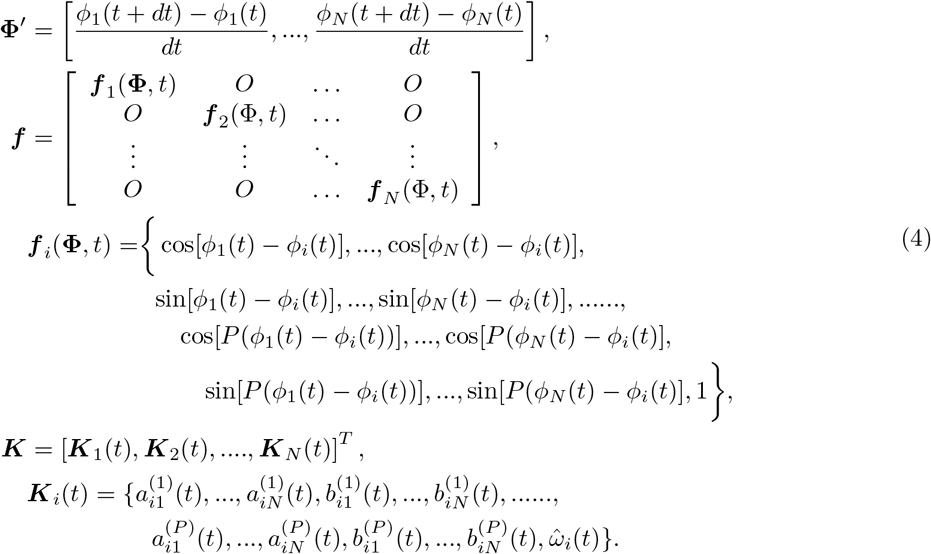

Note that **Φ′** indicates a phase velocity with a smaller time step *dt*. Moreover, ***K*** indicates the parameter vector of *P*-th Fourier coefficients. ***f*** indicates the design matrix, which is embedded in the vector ***f**_i_*(Φ, *t*) containing the cos and sin term of each *i*-th oscillator at time *t*. In the following numerical simulations (see section 2.5.1 for more details), we set the time step as *dt* = 0.01. Moreover, for the real measured data, the time step *dt* corresponds to the sampling interval of the data.

Given the prior distribution of the model parameters, the parameter ***K*** in Eq. (4) can be calculated using the Bayesian rule (Bishop, 2007; Sarris, 1973), as follows:

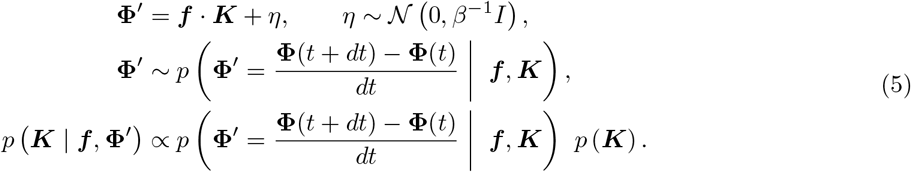

Here, in our proposed method, the prior distribution *p* (***K***) is assumed to have a multivariate normal distribution 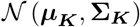, where ***μ_K_*** and **Σ_*K*_** indicate the mean and covariance of the distribution, respectively. The parameter *β* is a precision factor of the observation noise that was fixed as *β*^-1^ = 0.0001 for all analyses of this study.

The update rule of the prior distribution is derived as follows (Sarris, 1973):

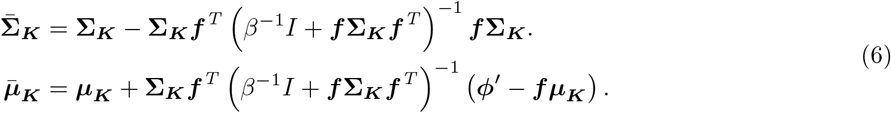

where 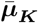 and 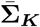 indicate the updated mean and covariance, respectively. After the update of the prior distribution, these parameters are set as 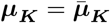 and 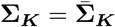 for the next updating process.

Moreover, the posterior predictive distribution of the model can be inferred using the following equation:

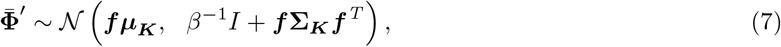

where 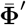 indicates the predicted phase velocity using the inferred model parameters. Based on the above procedure, the temporal changes in networks were estimated based on sequential Bayesian inference.

Our proposed method used the KL divergence between the prior and posterior distribution as an index of the change in dynamical structures, as shown in Eq. (8), and the value of this KL divergence is defined as the change point score (CPS) as follows:

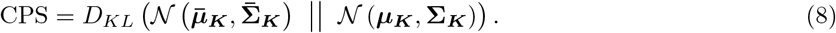

where ***μ**_K_* and 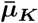 indicate the mean of the parameter *K* for the prior and posterior distribution, respectively. Also, **Σ**_*K*_ and 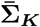 indicate the covariance of the parameter *K* for the prior and posterior distribution, respectively. Using the above process, we can detect change points in network dynamics while sequentially estimating dynamical structures in the brain using the KL divergence value.

In summary, our proposed method consists of the following seven steps.

1. Give the initial value of **μ_*K*_** and **Σ_*K*_** for the prior distribution of the model parameter
2. Get new single samples for each phase **Φ**(*t*), and calculate new variable ***f*** and **Φ**′(*t*)
3. Update predictive distribution of the model
4. Update the prior’s parameters
5. Calculate the KL divergence between prior and posterior model parameters (i.e., evaluate the change point score)
6. Substitute parameters as 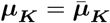 and 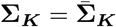 for the next Prediction step
7. Repeat steps 2 to 6

According to the above-mentioned description, the model parameter updating process is conducted on a sample-by-sample basis, and the metric of change point score using the KL divergence is also calculated after updating the prior of the model parameters sample-by-sample. The Python script of our proposed method is provided in the following GitHub repository: https://github.com/myGit-YokoyamaHiroshi/ChangeDetectSim

### 2.3. Evaluations of network estimation accuracy

In the following numerical simulations (see section 2.5.1), for each iteration of the model updating steps, we evaluated the error of the structure of network couplings with the mean absolute error (MAE) at time *t* to evaluate the estimation accuracy, as follows:

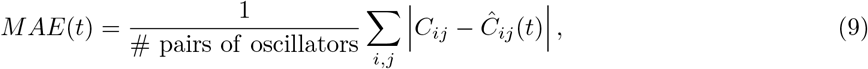

where *C_ij_* indicates the exact coupling strength value between the pair of oscillators *i*, *j* in the synthetic data, and *Ĉ_ij_*(*t*) indicates the estimated value of coupling strength between the *i*-th and *j*-th oscillators at time *t*. The value of *Ĉ_ij_*(*t*) at time t was calculated using estimated Fourier coefficients *â_ij_*(*t*) and 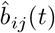 predicted by sequential Bayesian inference.

### 2.4. Evaluations of change detection accuracy and threshold selection

In most cases of change point detection problems, the frequency of anomaly changes was lower than the normal state frequency and such changes emerged suddenly. Therefore, to account for a minor delay in the change detection, we defined the “error tolerance area” after the true time-point of the changes (Gensler & Sick, 2014; Kovécs et al., 2020), as shown in Figure 2. The time samples corresponding to the change point scores higher than or equal to the pre-defined threshold *ζ* were labeled as positive; otherwise, the time samples were labeled as negative. Then, labels were categorized, as shown in Figure 2A–D. If positive labels were within the error tolerance area, the samples from the first positive point to the end of error tolerance area were tagged as True Positive (Figure 2A). By contrast, if no positive label was observed within the error tolerance area, all of the time-points corresponding to this area were tagged as False Negative (Figure 2D). For time-points outside the error tolerance area, time-points with no detection were tagged as True Negative (Figure 2B), and time samples with detection were tagged as False Positive (Figure 2 C). After categorizing the labels, the true positive rate (TPR) and the false positive rate (FPR) were calculated using the following equations:

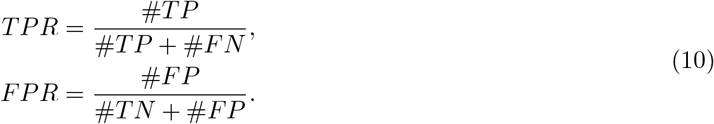

**Figure 2:**
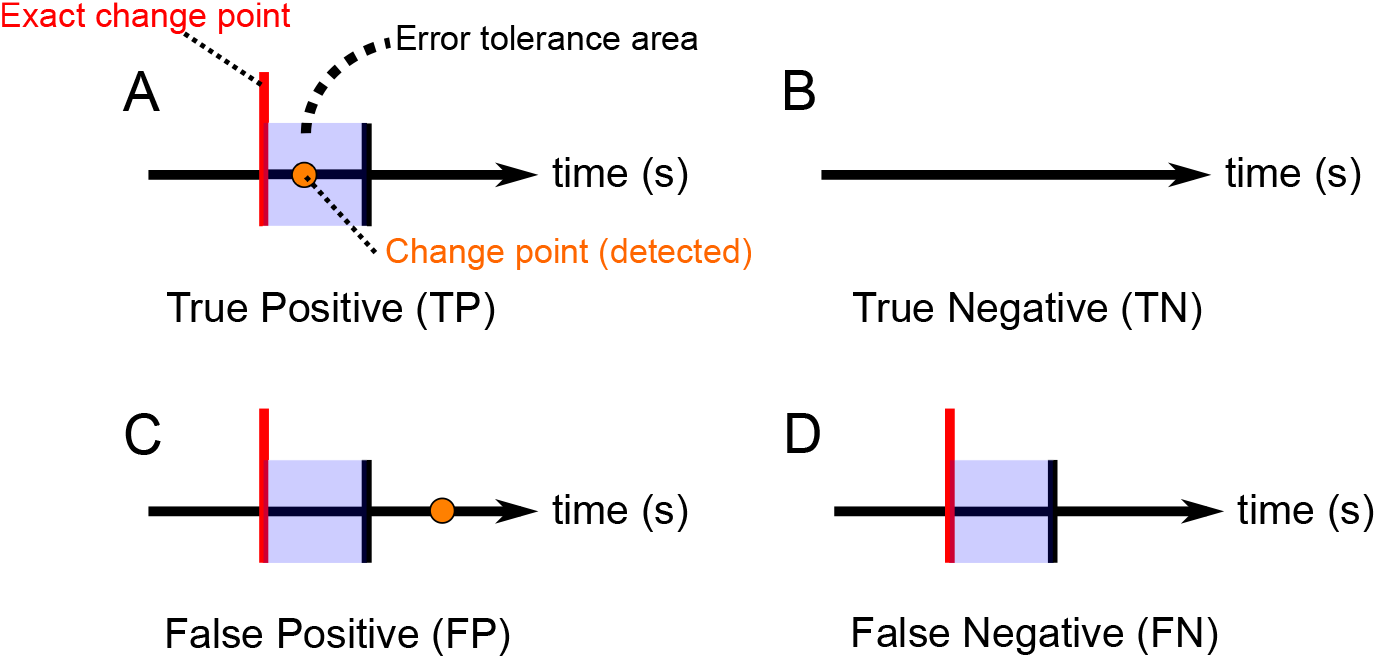
Schematic depiction of the change point evaluation. (A-D) Relevant situations for the evaluation of the detected change points. The red vertical lines indicate the change point. The blue shaded areas and the black vertical lines indicate the error tolerance area. If the change point was detected within the error tolerance area, the sample from the first significant point to the end of this area were labeled as True Positive (A).

To evaluate the detection accuracy, the receiver operating characteristic (ROC) curve and the area under the ROC curve (AUC) were calculated. To plot the ROC curve, the threshold *ζ* was changed from 0 to *ζ_max_*, and TPR and FPR were then calculated for each value of the threshold (Liu et al., 2013). After drawing the ROC curve, the AUC value was calculated as the accuracy of change point detection. Moreover, an optimal threshold *ζ*\* was also evaluated through the ROC curve analysis. In our study, an optimal threshold *ζ*\* was selected between 0 and *ζ_max_* to ensure that the following index was maximized:

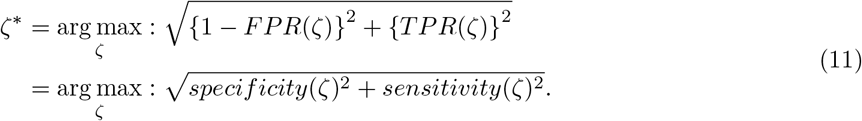

### 2.5. Validation of the proposed method

#### 2.5.1. Numerical simulation

In this study, we conducted three different simulations. The first simulation was conducted to confirm the number of network oscillators and the number of model learning iterations required to guarantee an accurate estimation of the network couplings when using our proposed method. As a short summary of the first simulation, we found that our method could be applied in the case where the number of oscillators *N_osci_* is less than or equal to 10 to guarantee the accurate network coupling estimation within 500 iterations of parameter updates (details of the results are provided in the Supplementary Materials). Considering the results of the first simulation, the second and third simulations were conducted to clarify whether our method can sequentially detect changes in the network couplings, using synthetic data generated by models with three or ten coupled oscillators, respectively. In the main manuscript, we only describe the results of the second simulation, and other the simulation’s results are provided in the Supplementary Materials.

The sample code of all numerical simulation described in this paper (both in the main manuscript and Supplementary Materials) is available at Github: https://github.com/myGit-YokoyamaHiroshi/ChangeDetectSim

##### 2.5.1.1. Procedure of the simulation

In this simulation, we applied our proposed method to the synthetic data generated by a phase-coupled oscillator model with three oscillators to clarify whether our method can sequentially detect changes in the network couplings. Moreover, to evaluate the effects of the sample size for the change detection accuracy and model parameter estimations, the four datasets, which contained different sample lengths of time-series as *N_t_* = [1, 500, 3, 000, 4, 500, 6, 000], were generated. In these datasets, to only focus on the effects of the sample size, other simulation conditions besides that of the sample size to generate the synthetic data were kept consistent.

The synthetic time series for each dataset was generated based on the stochastic phase-coupled oscillator model approximated with a first-order Fourier series using the numerical integration with the Euler-Maruyama method (see the Supplemental Materials for a more detailed description of the procedure of synthetic data generation). Note that the time step dt to solve the differential equation of the stochastic phase-coupled oscillator model was set as *dt* = 0.01 (i.e., the total time length of each dataset was set as *t* = 15, 30, 45, and 60s). Moreover, when generating for each dataset, the same initial value of phase was selected as Φ(0) = {*ϕ*_1_(0), *ϕ*_2_(0), *ϕ*_3_(0)} = {0, *π*, *π*/2}. The generated synthetic time series consisted of three segments separated by two events for each dataset. The time intervals of each segment were according to the sample length conditions, as shown in Table 1.

**Table 1:** Time interval for each segment. Four data sets of the synthetic data with different sample length were generated using the parameter setting shown in Figure 3. The synthetic time series consisted of three segments separated by two events, at distinct time depending on the sample length condition for each dataset.

| 1 ) $N_t = 1,500$ (15s) | | 2 ) $N_t = 3,000$ (30s) | |
| --- | --- | --- | --- |
| segment 1 | $0 \leq t(s) \leq 5$ | segment 1 | $0 \leq t(s) \leq 10$ |
| segment 2 | $5 \leq t(s) \leq 10$ | segment 2 | $10 \leq t(s) \leq 20$ |
| segment 3 | $10 \leq t(s) \leq 15$ | segment 3 | $20 \leq t(s) \leq 30$ |
| 3 ) $N_t = 4,500$ (45s) | | 4 ) $N_t = 6,000$ (60s) | |
| segment 1 | $0 \leq t(s) \leq 15$ | segment 1 | $0 \leq t(s) \leq 20$ |
| segment 2 | $15 \leq t(s) \leq 30$ | segment 2 | $20 \leq t(s) \leq 40$ |
| segment 3 | $30 \leq t(s) \leq 45$ | segment 3 | $40 \leq t(s) \leq 60$ |

In the first event, the Fourier coefficients *a_ij_* and *b_ij_* changed, which indicated the occurrence of changes in the dynamical structure of networks in the boundary between segments 1 and 2 (hereafter, this event is called “structural change”). In the second event, the Fourier coefficients *a_ij_* and *b_ij_* did not change between segments 2 and 3; however, the observation noise, which does not directly contribute to temporal dynamics, drastically changed (hereafter, this event is called “noise-level change”). The exact values of the parameters *a_ij_* and *b_ij_* were set as shown in Figure 3. The natural frequency 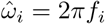 of oscillator *i* was set at a fixed value with *f* = {*f*_1_, *f*_2_, *f*_3_} = {4.31, 4.48, 3.70}. Moreover, the noise-scaling factor *p_i_* for calculating numerical integration using the Euler-Maruyama method (see the Supplementary Materials for more details about the noise scaling factor) was set as Eq. 12.

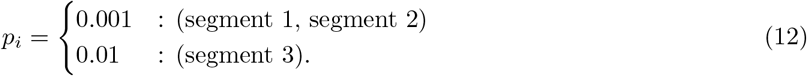

**Figure 3:**
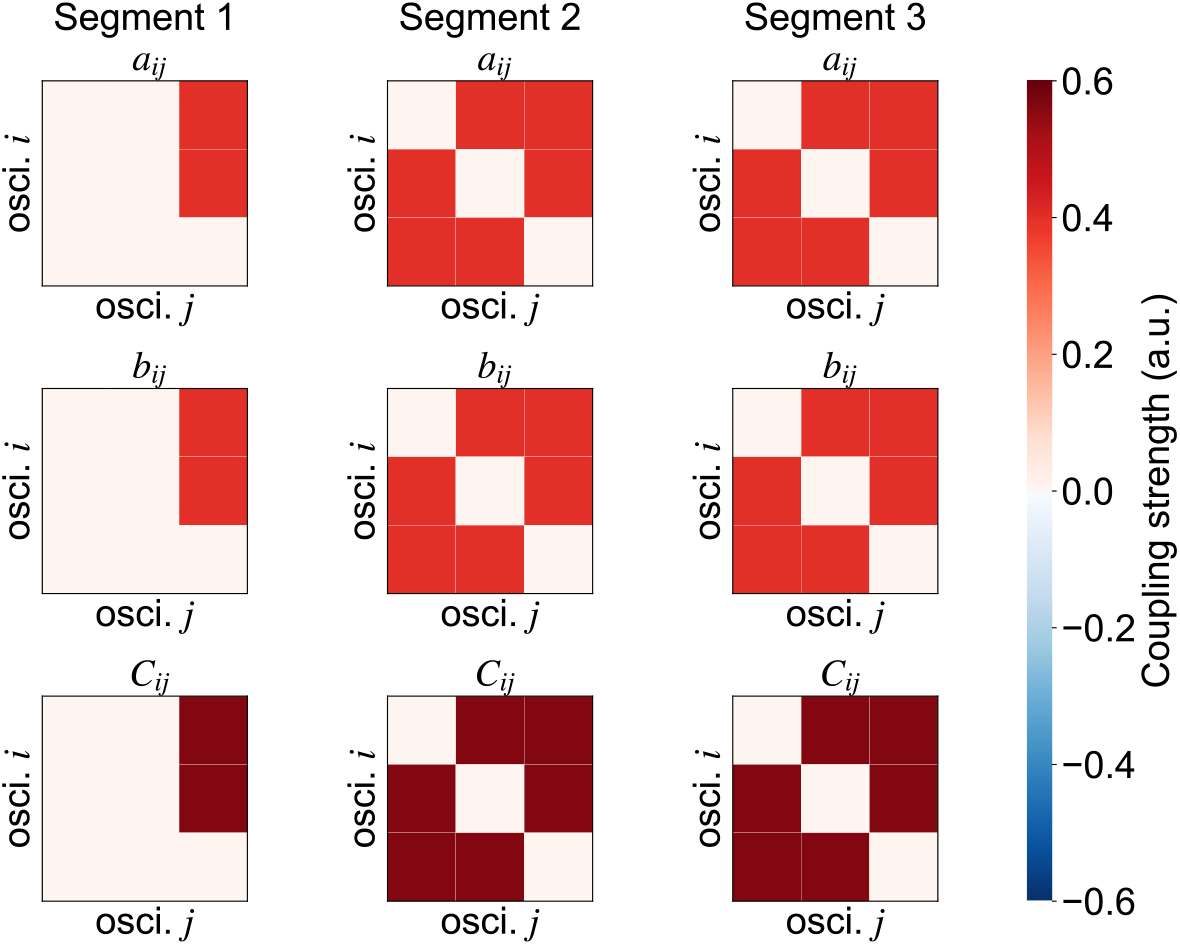
Parameter settings in numerical simulation. True parameter settings in each segment. The time series of synthetic data was generated using the function Γ_*ij*_ (*ϕ_j_* – *ϕ_i_*) approximated by the first-order Fourier series using these parameter settings in each segment. *a_ij_* and *b_ij_* indicate the Fourier coefficient between the *i*-th and *j*-th oscillators. *C_ij_* is the network couplings between the *i*-th and *j*-th oscillators.

After generating the synthetic data with the above-mentioned parameter settings, our method was applied to each dataset of the synthetic data. The estimation performance of change detection and network couplings were evaluated with the performance metrics. In this numerical simulation, to draw the ROC curve, the values of *ζ* were selected from 0 to *ζ_max_*, and TPR and FPR were then calculated for each threshold value. Note that the values of *ζ_max_* were set as the mean(KL) + 5SD(KL), where the mean(KL) and SD(KL) indicate the mean and standard deviation of the KL divergence (i.e., change point score) across samples. Furthermore, in the ROC curve analysis for the numerical simulations, to strictly validate the detection accuracy of our proposed method, we did not consider the error tolerance area after the true change point (i.e., the sample length of the error tolerance area for change point detection was set as zero for this numerical simulation).

##### 2.5.1.2. Comparison with the previous method

To compare the change detection performance of our method with that of the conventional method, the network changes in synthetic data were also evaluated using the adaptive vector auto-regressive (VAR) model with Kalman filter (Xiong & Zhou, 2013; Lie & van Mierlo, 2017). As mentioned in the Supplementary Materials, since a parameter estimation for linear regression using sequential Bayesian inference with the Gaussian prior is a special case of Kalman filter, we selected this adaptive VAR-based method for the comparison with the proposed method. In this method, the time-series were fitted with the following *P*-th order of VAR model:

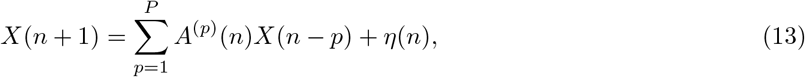

where *A*^(*p*)^(*n*) and *X*(*n*) indicate *N_osci_* × *N_osci_* matrix of model coefficients at time sample *n* and *N_osci_* × 1 vector of time-series with *p*-th delay from time sample *n*, respectively. The model coefficient *A*^(*p*)^(*n*) were recursively estimated from the observed data using the Kalman filter. Note that the model order *P* is a selected value where the maximum log-likelihood is within the range of the 1st to 20th order. Using its estimated parameter *A*^(*p*)^(*n*), we can also estimate causal relationship from the variables *i* to *j* using the partial directed coherence (PDC), as follows:

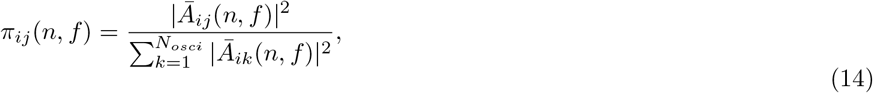

where,

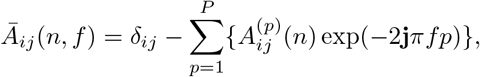

where *δ_ij_* = 1 whenever *i* = *j*, and otherwise *δ_ij_* = 0. *Ā_ij_*(*f*, *n*) is the Fourier-transformed *p*-th VAR coefficient 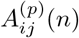 with frequency *f* at time sample *n*. 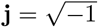 indicates the imaginary unit. In our study, the frequency-averaged value of PDC, such as 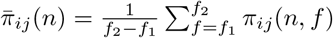 was applied as an index of network coupling with the VAR model at time sample *n*, and frequency band [*f*_1_, *f*_2_] was set as [*f*_1_, *f*_2_] = [3, 5] in this numerical simulation, since the natural frequency of the generated synthetic data was set to around 4 Hz. More details of the algorithm of parameter estimation based on the Kalman filter are reported elsewhere (Xiong & Zhou, 2013; Lie & van Mierlo, 2017). The change point score is defined as the square of Mahalanobis’ distance between the observed data and estimated data for each time sample (so-called Hotelling’s T-square metric) based on the following equation:

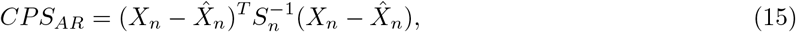

where *X_n_* and 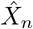 are the observed and estimated data at time sample *n*. *S_n_* is the prediction error covariance at time sample *n*, determined by the Kalman filter. By applying the VAR-based method, the accuracy of change point detection and network coupling estimation were also evaluated for each dataset of generated synthetic data in the same manner as our proposed method, and these results were compared with those obtained using our method. Moreover, when applying this VAR-based method for the synthetic phase data in this simulation, the time series *X* was defined as *X* = cos(Φ(*t*)), where Φ(*t*) = {*ϕ*_1_(*t*), *ϕ*_2_(*t*), *ϕ*_3_(*t*)} is the time-series data of the synthetic phase.

#### 2.5.2. Validation of our method using EEG data

Finally, to confirm its neurophysiological validity, we applied our method to real human EEG data obtained by McFadden et al. (2014). This dataset includes EEG recordings from healthy human participants who were passively listening to 40-Hz AM white noise. In general, when listening to 40-Hz AM white noise, 40-Hz oscillatory activity is recorded over the auditory areas (the so-called auditory steady-state response; ASSR (McFadden et al., 2014; Rauschecker & Scott, 2009; Reyes et al., 2004; Ross et al., 2005)). In addition, a recent ASSR study that used a directional connectivity analysis (Ying et al., 2015) suggested that network connectivity associated with 40-Hz ASSR in healthy humans is mainly distributed at central and fronto-temporal areas. We applied our method to the open EEG dataset to assess whether our method can detect the dynamical changes in neural activity induced by auditory stimuli, as reflected by EEG recordings.

##### 2.5.2.1. Open EEG dataset

In the dataset used, EEGs were recorded with a 10-10 system-based 64-channel system (Neuroscan, Inc., North Carolina, USA; sampling frequency: 1,000 Hz) in 19 healthy participants who were passively listening to 40-Hz AM noise (McFadden et al., 2014). The AM noise was binaurally presented through foam insert earphones at 75 dB SPL for 0.5 s with an inter-trial interval of 1.0 s (McFadden et al., 2014). The participants performed over 200 trials of this passive listening task (McFadden et al., 2014). A more detailed description of the task settings is provided in the original paper (McFadden et al., 2014). In the following validation of this study, we only used EEG data recorded during the 40-Hz AM noise in participant ID no. 1. Out of the recorded trials, 225 trials were applied and used for the change point detection analysis described in the following section.

##### 2.5.2.2. Preprocessing and change point analysis

EEG signals were re-referenced off-line to the average of the signals from left and right earlobe electrodes. Then, to reduce the effects of eye movement and eye blink signals, we applied an artifact removal method with empirical mode decomposition (Flandrin et al., 2004, 2005; Molla et al., 2010). A detailed description of this empirical mode decomposition-based electrooculographic artifact removal method is provided in previous articles (Flandrin et al., 2004, 2005). After removing the artifacts, EEGs were converted to current source density signals (Kayser et al., 2006). To extract phase oscillatory dynamics around 40 Hz, the current source density signals were band-pass filtered between 38 and 44 Hz with a zero-phase FIR filter and converted to analytic signals using the Hilbert transform. Next, phase angles were extracted from the analytic signals of the current source density signals and divided into 1.7 s epochs (−1.0 to 0.7 s, 0.0 s: stimulus onset) with a 0.7 s overlap between neighboring epochs.

After preprocessing, the following change point analyses were individually applied to the time series of EEG phase data for each epoch, in the same manner as in the numerical simulation. In the analyses, we used data recorded from 10 channels (F3, F4, C3, Cz, C4, P3, Pz, P4, O1, and O2); these 10 channels were selected because the condition for the number of oscillators needed to confirm that the accuracy was high enough within 500 iterations of parameter updates was *N_osci_* = 10 (see Supplementary Materials for more details). For each epoch, the first 1,000 samples of the pre-stimulus period (i.e., −1.0 ≤ *t* ≤ 0.0) were applied to the prior learning for the model, and the later 0.2 intervals of this period (i.e., −0.2 ≤ *t* ≤ 0.0) were defined as the baseline period to calculate the threshold *ζ* of the ROC curve analysis for the accuracy evaluation of change point score. In the EEG analysis, to draw the ROC curve, the value of *ζ* was selected from 0 to *ζ_max_*, and TPR and FPR were then calculated for each value of the threshold. Note that the value of *ζ_max_* was set as the mean(KL_base_) + 5SD(KL_base_), where the mean (KL_base_) and SD(KL_base_) indicate the mean and standard deviation of the KL divergence (i.e., change point score) in baseline periods. Using this threshold, the ROC curve and AUC were calculated in same manner using the numerical simulation for each single epoch individually. Moreover, in this evaluation for EEG data analysis, to account for a minor delay in the neuronal response, we defined the “error tolerance area” as 0.1 s after the true time-point of the stimulus onset/offset for evaluation of the TPR and FPR.

##### 2.5.2.3. Validations of the change detection accuracy

To evaluate the statistical significance of the detection accuracy, the median values of each index over all trials were calculated, and the significance of these values was confirmed using a one-tailed permutation test. The null distribution for this permutation test was generated using the following steps:

1. The significance of the change point score of each time sample was assessed based on the threshold *ζ*.
2. If the score was greater than or equal to the *ζ*, the time samples were labeled as positive. Otherwise, the samples were labeled as negative.
3. The labels (positive or negative) were shuffled.
4. The TPR and FPR were evaluated according to these randomized labels.
5. Steps 1 to 3 were repetitively executed between 0 ≤ *ζ* ≤ mean(KL_base_) + 5SD(KL_base_) to create the randomized ROC curve.
6. Steps 1 to 4 were carried out for each trial, and the median of the randomized ROC curves for all trials was calculated.
7. All steps were repetitively executed 1,000 times, and the null distribution of the ROC was generated from those 1,000 randomized median ROC curves.

For this statistical analysis, the median of the original ROC curves of all epochs were compared with the null distribution using the above-mentioned steps.

## 3. Results

### 3.1. Numerical simulation

As mentioned in the section 2.5.1, considering the results of supplementary simulation described in the Supplementary Materials, we applied our method to ≤10 oscillators in this study. We first conducted numerical simulations using four datasets, which contain the synthetic data generated by a phase-coupled oscillator model of three oscillators with different sample length conditions. Both our proposed method and the VAR-based method were applied to these datasets, and we compared the change point detection accuracy between the two methods. The results of this simulation are shown in Figures 4, 5, and 6.

**Figure 4:**
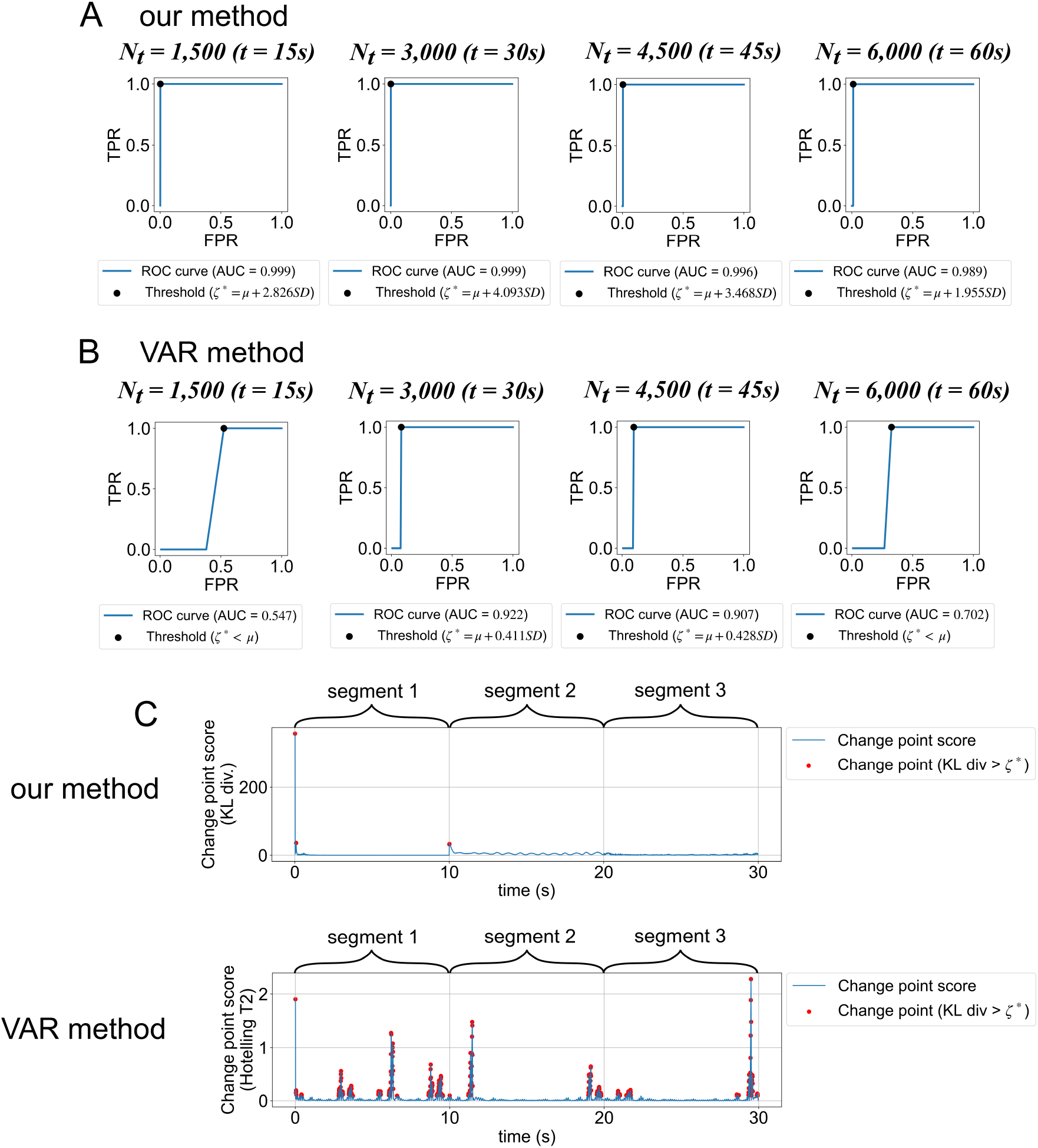
Change point detection in the numerical simulation. (A and B) Results of the ROC curve analysis for each dataset based on the proposed method and the VAR-based method, respectively. Blue lines indicate the evaluated ROC curve with a threshold 0 ≤ *ζ* ≤ *μ* + 5*SD*, where *μ* and *SD* indicate the mean and standard deviation of the change point score across samples for each change detection method. Black dots indicate the location of an optimal threshold *ζ*^*^. The AUC score and selected value of *ζ*^*^ are shown in each figure legend in the form of “*μ* + *N* · *SD*”. If the selected value of *ζ*\* is smaller than *μ*, this is presented as “*ζ*\* < *μ*”. (C) Example of the time-course of the estimated change point score and selected time samples of the change point with the optimal threshold *ζ*^*^ in the dataset *N_t_* = 3, 000 (*t* = 30s). The upper and lower panels indicate the time course of the change point scores according to the proposed method and the VAR-based method, respectively. The red dots indicate the selected change points with the threshold *ζ*\*.

**Figure 5:**
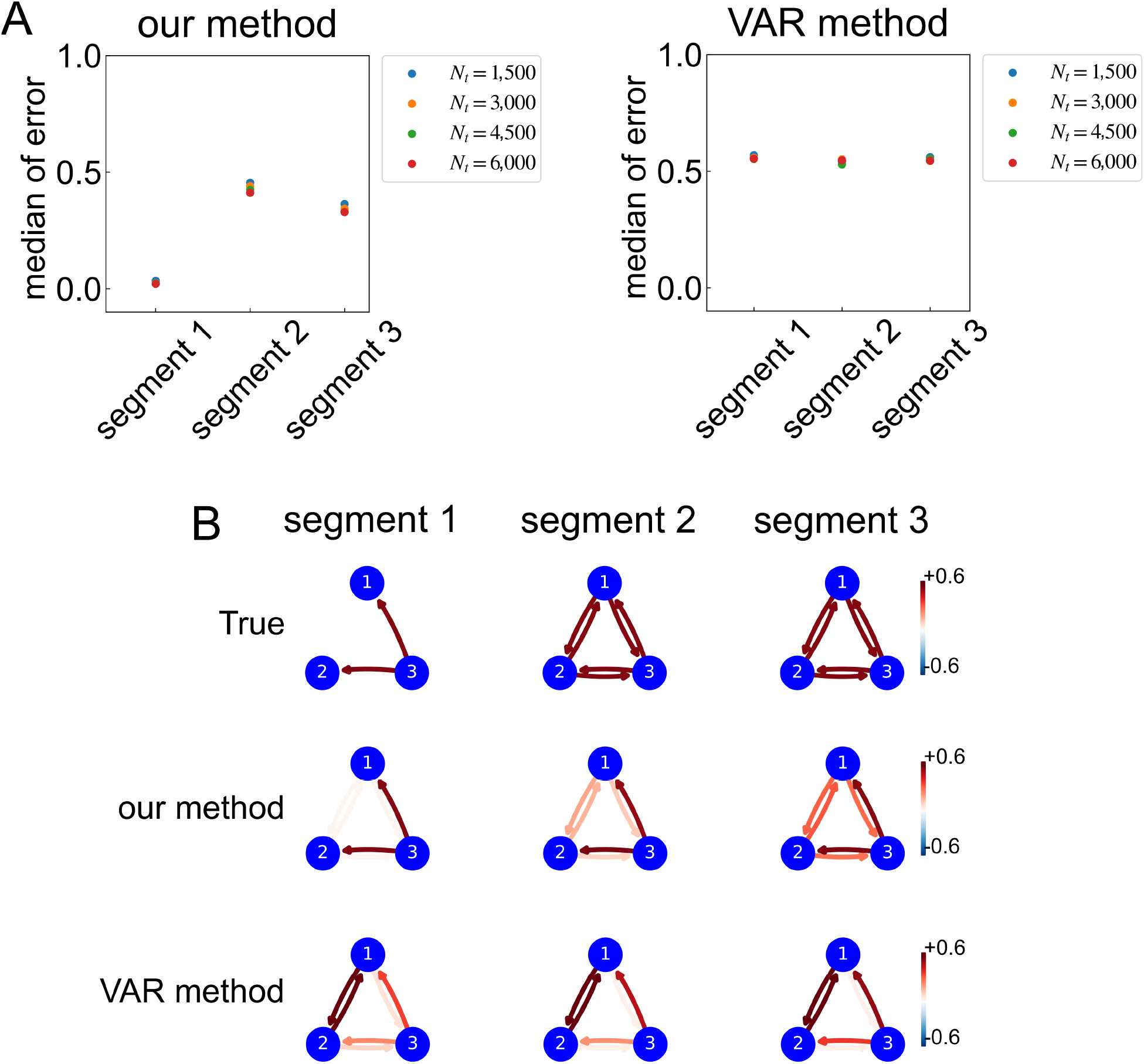
Comparisons of the accuracy of the network couplings estimation between the two estimation methods. (A) Comparison of the medians of the mean absolute error (MAE) of the network couplings between the three segments. Colored dots indicate the median of the MAE of four datasets with distinct numbers of samples. (B) Comparisons of true and predicted network structures based on two methods in each segment. The upper three images show the true network couplings for each segment. The middle and lower three images show the estimated network couplings based on the proposed method and the VAR-based method in the datasets: *N_t_* = 3, 000, respectively.

**Figure 6:**
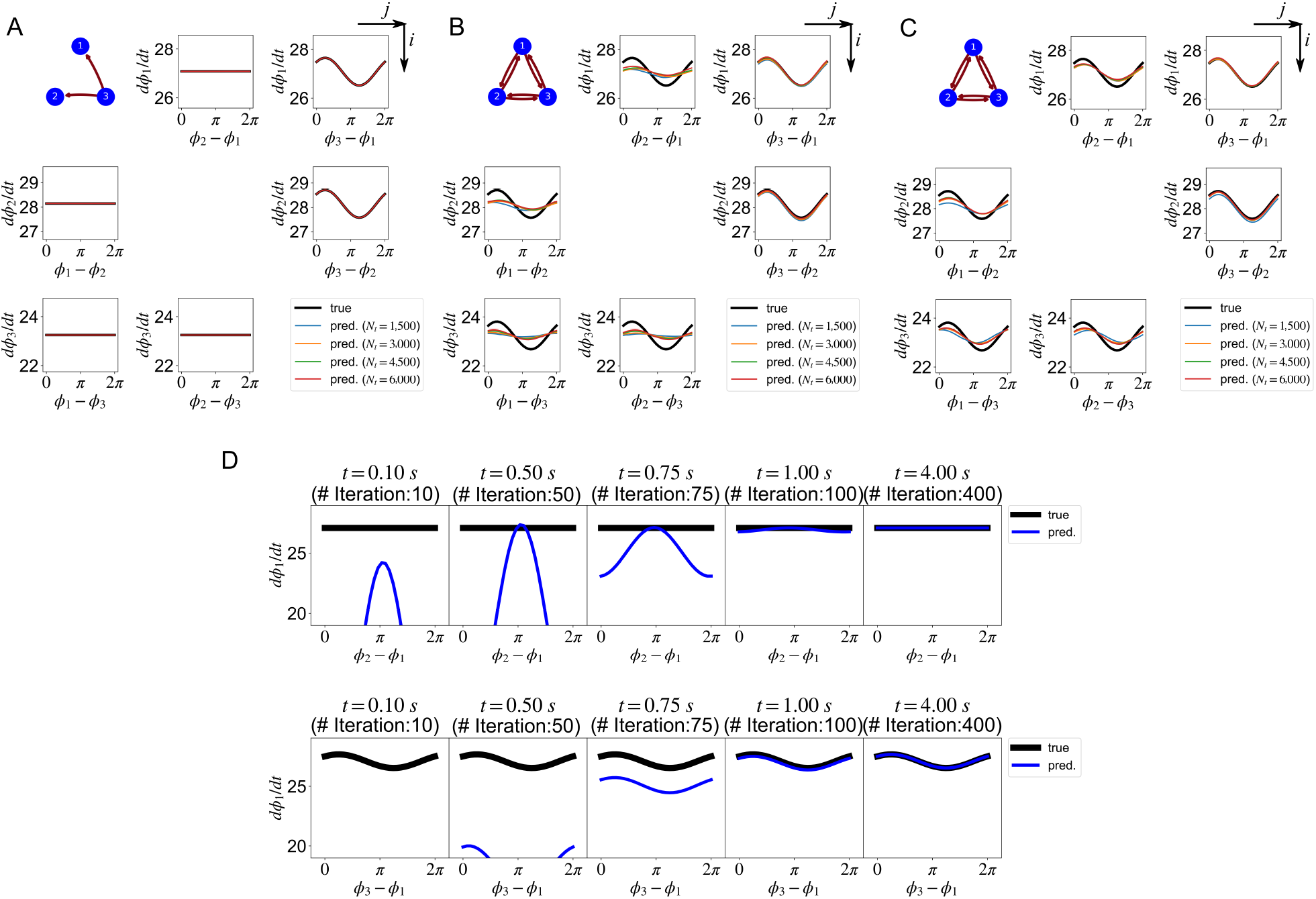
Estimated functions Γ_*ij*_ (*ϕ_j_*(*t*) – *ϕ_i_*(*t*)) from synthetic data in numerical simulation. (A-C) Results for segments 1, 2, and 3, respectively. The upper left diagram in each panel shows the actual network coupling structure of the synthetic data. The black bold lines show the actual phase interaction curves for all pairs of oscillators. The colored lines show the median of estimated phase interaction functions for each simulation dataset. The shape of curves for the interaction functions indicates an interaction from the *j*-th to *i*-th oscillators, estimated by our proposed method in each segment. The phase interaction function would be flat in the absence of an interaction between the *i* and *j*-th oscillators. (D) Typical examples of the estimated function Γ_*ij*_ (*ϕ_j_*(*t*) – *ϕ_i_*(*t*)) of each parameter updating iteration (dataset: *N_t_* = 3, 000, segment 1). The upper panels show the results of the changes in the estimated interaction function from the second to the first oscillators for each iteration. The lower panel show the results of the changes in the estimated interaction functions from the third to the first oscillators for each iteration.

First, we compared the results of ROC curve analysis for each method. Figure 4A and B show the ROC curves and AUC score for each dataset; the AUC score, which was calculated according to the evaluated ROC curves, was, overall, higher in our method than in the VAR-based method. Moreover, the shape of the ROC curve and its AUC score estimated using the VAR method are affected by the difference of the time-samples, whereby our method consistently estimated a higher change detection accuracy (*AUC* >= 0.9989) without dependence on the sample size. Furthermore, as shown in Figure 4C, while the VAR method has a high false detection rate of change points, our method can correctly detect the change point in the boundary of segments 1 and 2. In addition, what the change point were not detected in around the boundary between segments 2 and 3 in our method is also important advantage of our method. This result indicates that our proposed method can correctly detect the change point exclusively for structural changes in the network couplings, without being influenced by observation noise. In other words, the results indicate that our method guarantees accurate change point detection of the network couplings with a significant noise tolerance. Figure 4C shows the time course of the change point score for the dataset of *N_t_* = 3, 000 results only; however, this tendency was consistently shown for the other datasets (*N_t_* = 1, 500, 4, 500, and 6, 000), as shown in the figure. A common tendency between our method and VAR-based method was that the first few time samples were falsely detected as change points, and so the model estimation was unstable for first few samples (see the interval of segment 1 in Figure 4C). However, even if such misdetection occurs in the early stage of model parameter learning, the accurate change point detection performance, without dependence on the sample size, is a clear advantage of our method.

We also compared the prediction accuracy of the network coupling between our method and the VAR-based method. Figure 5B shows the comparison between the true and predicted network for each segment. In our method, the predicted network couplings were calculated as the median coupling strengths determined by Eq. (3) with estimated Fourier coefficients for each segment. In the same way, for the VAR-based method, the predicted network couplings were calculated as the median coupling strengths determined by PDC with estimated AR-coefficients for each segment. As shown in Figure 5B, while exact and estimated networks using our method were very similar for each segment, the estimated networks in the VAR-based method were falsely detected for each segment. In particular, our method was able to detect network coupling correctly in segment 1, even though this is where indirect network coupling between the 1st and 2nd oscillators, caused by a common drive from the 3rd oscillator as a confounder, can easily occur. In contrast to the results of our method, the VAR-based method falsely detected the coupling between the 1st and 2nd oscillators. In addition, as shown in Figure 5A, the estimation error of each segment was higher in the VAR-based method than in our method for each sample size condition.

To further examine whether our proposed method could detect the time-evolving dynamics of oscillatory networks, we compared the predicted function Γ_*ij*_ (*ϕ_j_*(*t*) – *ϕ_i_*(*t*)) with the actual interaction function of each pair of oscillators for each segment for all datasets. Figure 6 shows the median of the functions Γ_*ij*_ (*ϕ_j_*(*t*) – *ϕ_i_*(*t*)) for each segment. The function Γ_*ij*_ (*ϕ_j_*(*t*) – *ϕ_i_*(*t*)) for each dataset was evaluated estimated Fourier coefficients for all time-points *t*, and the median of these estimated functions were then calculated over the time samples for each segment. If there is no interaction between the *i* and *j*-th oscillators, the function Γ_*ij*_(*ϕ_j_*(*t*) – *ϕ_i_*(*t*)) will be flat, indicating no interaction between the two oscillators. As shown in Figure 6A, our model made estimations such that the function Γ_*ij*_ (*ϕ_j_*(*t*) – *ϕ_i_*(*t*)) would be flat in the case of no interaction between the pair of oscillators without dependence on the time-sample conditions. Moreover, as shown in Figure 6B and 6C, the predicted functions Γ_*ij*_ (*ϕ_j_*(*t*) – *ϕ_i_*(*t*)) were similar between segments 2 and 3, even though the noise level and sample lengths were different. This indicates that our method can capture changes in the phase interaction function as well as in network coupling.

Furthermore, to confirm the detailed profiles of the changes in the predicted Γ_*ij*_(*ϕ_j_*(*t*) – *ϕ_i_*(*t*)) with the number of parameters updating iterations, the predicted Γ_*ij*_(*ϕ_j_*(*t*) – *ϕ_i_*(*t*)) in segment 1 were plotted with several iteration numbers. Figure 6D shows the results for the function Γ_12_ (*ϕ*_2_(*t*) – *ϕ*_1_(*t*)) and Γ_13_ (*ϕ*_3_(*t*) – *ϕ*_1_(*t*)) in the time intervals between 0 to 4 s in the datasets: *N_t_* = 3,000. As shown in the figure, the predicted functions Γ_12_(*ϕ*_2_(*t*) – *ϕ*_1_(*t*)) and Γ_13_ (*ϕ*_3_(*t*) – *ϕ*_1_(*t*)) converged to the shape of the true interaction function within 100 iterations of the parameter updating, and this tendency was consistently shown for each pair of oscillators, regardless of the sample size conditions of the datasets.

### 3.2. Scalp EEG data

Finally, we evaluated the neurophysiological validity of our method by applying it to EEG data that were shared by McFadden et al. (2014). Figure 7A–C show the typical correct (Figure 7A, B) and incorrect results (Figure 7C) of change point detection in EEG data from participant 1. The left panel in Figure 7A–C shows the evaluated ROC curve on a single-trial basis with the threshold *ζ* between 0 and mean (KL_base_) + 5SD(KL_base_). The right-hand panels in Figure 7A-C show the change point detection results with threshold *ζ*\* evaluated by ROC curve analysis. As shown in these figures, the change point scores were significantly increased around the stimulus onset and offset in epochs for which the change point within the error tolerance area was successfully detected, as shown in Figure 7 A and B, while in epochs for which there was an incorrect detection, the significant increase of the change point scores was outside the error tolerance area (Figure 7C). To confirm the tendency over all epochs in more detail, the median of the original ROC curves over 255 trials was calculated. As can be seen in Figure 7D, the AUC score, which was calculated according to the median of the original ROC curves, was 0.824, which was significantly larger than the AUC score evaluated with the median of permutation ROC curves. Therefore, although the ability of our method to ensure the validity of change point detection was limited by the number of EEG electrodes, the results indicate that our method successfully detected the changes in EEG phase oscillatory dynamics with significant accuracy.

**Figure 7:**
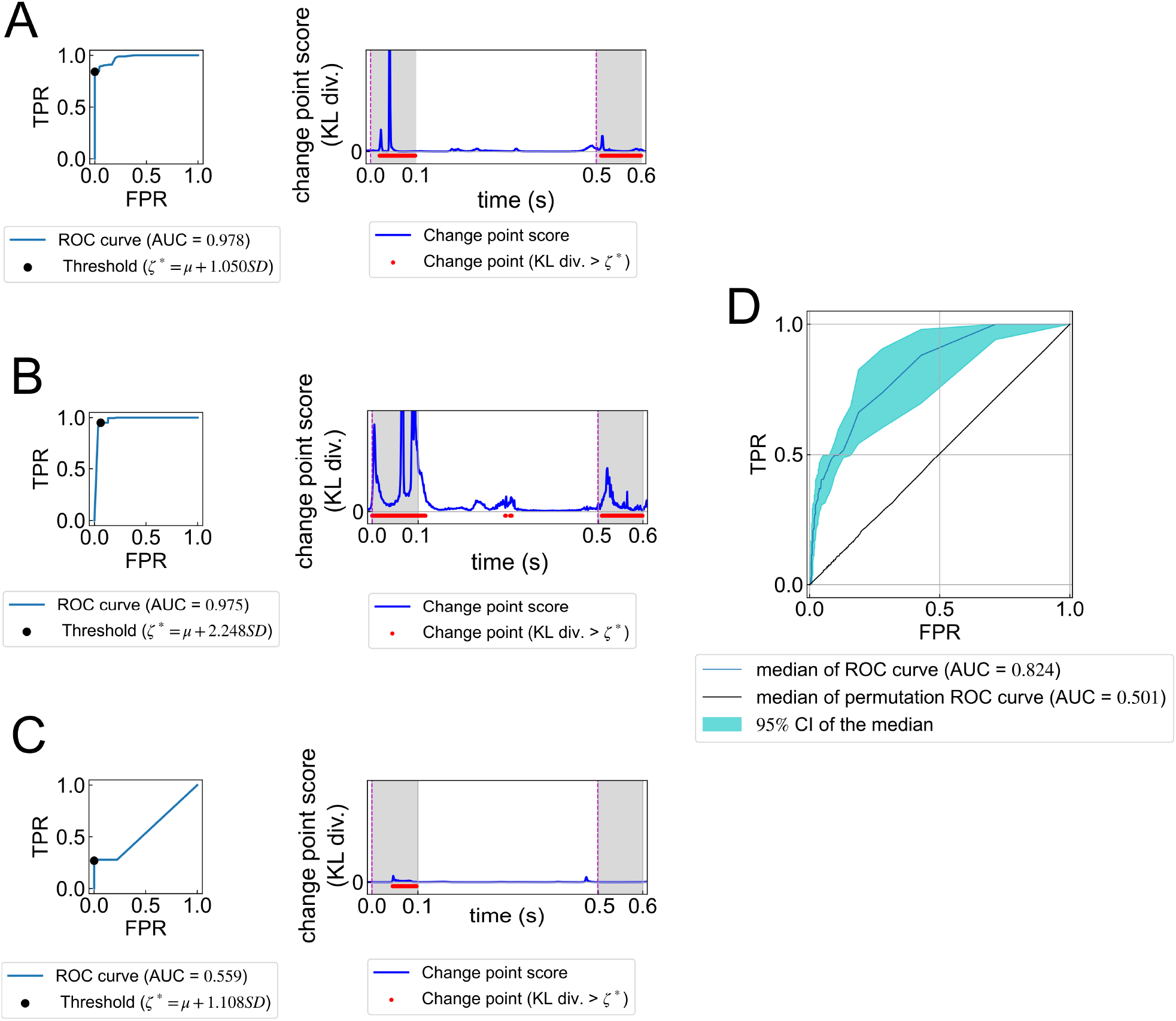
Change point detection analysis in the EEG dataset. Typical examples of correct epochs in change point detection in EEG data (the optimal threshold *ζ*^*^ was individually selected by the ROC curve analysis for each epoch). The left and right panels show the results of the ROC curve analysis and the time-course of change point scores, respectively. In the right-hand panels of (A) and (B), the pink dashed line and gray shaded area indicate the stimulus onset/offset and the error tolerance area, respectively. (C) A typical example of incorrect epochs in change point detection. This result is presented in the same way as in (A) and (B). (D) The median ROC curve of all 255 epochs. Blue lines and the light blue shaded area indicate the median ROC curve and 95% confidence intervals of the median of ROC curves. Black lines indicate the permutation median of the ROC curves. The value of the AUC score shown in (D) was calculated based on the median ROC curve of all 255 epochs.

## 4. Discussion

In this study, we proposed a new approach to detect the change points of brain networks by extending the data-driven method to estimate phase oscillatory dynamics based on sequential Bayesian inference. To validate our proposed method, we applied it to numerical and empirical neurophysiological data. The numerical data analyses confirmed that the change point score of our method significantly increases only when structural changes in the dynamical network occur, and not when noise-level changes in the dynamical network occur. These results support the two following assumptions for our proposed method: (1) time-evolving dynamics and data structures of functional brain networks can be represented by a phase-coupled oscillator model, and (2) temporal changes in dynamical brain structures are reflected in changes in model parameters. Furthermore, the neurophysiological data analysis revealed that our method was able to detect the change point of phase oscillatory dynamics induced by auditory stimuli with an *AUC* = 0.824 accuracy, which was based on the median of original ROC curves for all epochs. These findings suggest that our proposed method has mathematical and neurophysiological validity. In the following section, we will discuss the neuroscientific significance of our proposed method, and will compare our results with those of previous studies that have employed time-varying functional connectivity analyses.

### 4.1. Advantages of our proposed method

The originality of our proposed method lies in the application of sequential Bayesian inference to estimate the phase interaction function of dynamical networks and to detect the change point of such dynamics. Using this approach, the changes in the network dynamics of the brain could be expressed as the changes in the model statistics regarding the parameters of the phase interaction function (Figure 1). Therefore, our method is not only able to visualize time-varying network couplings of the brain, but can also represent the extent of the time-varying changes in dynamical brain structures using an information theoretical index (the KL divergence).

Relevant previous studies have reported two typical methods that can be used to estimate temporal changes in the network connectivity of neural activity. First, relevant conventional studies to estimate time-varying changes in functional connectivity of the human brain (Hutchison et al., 2013; Damaraju et al., 2014). This method extends the functional connectivity analysis to estimate time-varying network structures by applying the sliding window method (so-called dynamic functional connectivity analysis; dFC analysis) (Hutchison et al., 2013; Damaraju et al., 2014). Although dFC analysis focuses on the temporal changes in brain networks, this method separately evaluates functional networks with static indices, such as the correlation between time-series data, for each time window. Thus, these analyses do not explicitly consider the time-evolving dynamics of time-series data, because multivariate time series with non-stationary dynamics usually contain temporal dependence and auto-correlation, as in a dynamical system. Moreover, in most of the studies using dFC, when evaluating the network structure in each time window, the coupling strength for each pair of brain areas is calculated based on the temporal correlation with the time-series data. According to prior evidence (Granger et al., 1974), correlation coefficients between two different time-series data tend to be large, regardless of the existence of causality between two time-series data. Therefore, this method could lead easily to the detection of incorrect network connectivity based on spurious correlations. The second method is the change point detection algorithm of the time-varying gene regulatory networks by combining the auto-regressive (AR) model and Kalman filtering (Xiong & Zhou, 2013). The Kalman filter is based on the state-space model framework with a Bayesian parameter updating approach. Xiong & Zhou (2013) combined the recursive parameter estimation of the AR model and Kalman filtering by giving the model parameters of the AR model for the state variables (i.e., model parameters in the AR model are estimated as state variables using Kalman filtering). Unlike the dFC approach, this method is more advanced in its ability to estimate time-varying network connectivity by considering the time-evolving dynamics based on the state-space model. As stated in the Introduction section, the method combining the auto-regressive (AR) model and Kalman filtering are also applied in neuroscience research in the model-based dFC analysis (Astolfi et al., 2008; Leistritz et al., 2013). Based on prior studies, in the numerical simulation of the current study, we also evaluated the change detection performance using this VAR-based method, as the prior method for change detection for dFC.

The VAR-based method combined with the Kalman filter is similar to our own method, in that it also applies a Bayesian approach to estimate the model parameters. However, this VAR-based method explicitly assumes that both the network interaction and temporal dynamics in the time series data are linear. In the case of synchronous neural oscillatory data, time-evolving dynamics driven by the interactions for each oscillator could be considered as a coupled nonlinear system. Considering the possibility that a nonlinear dynamical system, such as a phase-coupled oscillator, underlies neural data might be important to estimate the time-varying network coupling for neural data. As shown in the comparison between our method and the VAR-based method (Figures 4 and 5), although the VAR-based method has a high false detection rate of change points, our method can correctly detect the change point only when structural changes in the network couplings occur. This suggests that making the assumption that a dynamical system underlies neural data could be an essential way to consider both temporal dependence and network coupling to deal with the change detection problem for neural time-series data.

Our proposed method advances the above-mentioned conventional methods (Hutchison et al., 2013; Astolfi et al., 2008; Leistritz et al., 2013; Xiong & Zhou, 2013) in the two following ways. First, our method considers both time-evolving dynamics and a nonlinear dynamical structure of functional brain networks. Unlike the dFC frameworks, our method adequately considers nonlinear and time-evolving properties of the brain activity by applying a dynamical model-based approach to estimate time-varying brain networks. The advantage of such a dynamical model-based approach is supported by the finding that misinterpretation caused by spurious correlations is largely reduced by applying a network system with nonlinear temporal dynamics. As can be seen in Figures 4 and 5, our method was able to correctly detect the network coupling and its changes, even in situations where the algorithm with a model-based method using linear model (VAR model) led to incorrect estimations. Second, our method can sequentially detect changes in dynamical brain structures based on Bayesian inference and an information-theoretic metric (i.e., KL divergence). Moreover, our method can consequently consider the temporal dynamics of time-series data by applying sequential Bayesian inference to estimate the time-varying changes in the model parameters. As a subsidiary advantage of applying the phase-coupled oscillator model to estimate network dynamics of the brain, directional network couplings could be detected, which is not the case with dFC analysis.

In summary, by adopting the property of time-evolving dynamics in the observed data using a dynamical model, the proposed method succeeded in providing a quantitative interpretation of the change point of time-evolving dynamics in brain networks using probabilistic inference and an information-theoretic metric.

### 4.2. Neuroscientific interpretation and generalizability

In this study, by supposing that the phase-coupled oscillator model is a representation of the time-evolving dynamics of functional brain networks, we expected the changes of functional brain networks to be quantified as changes in dynamical structures of the model parameters. Based on this assumption, we applied our proposed method to a multi-channel EEG time series that was measured from a healthy participant who was passively listening to 40-Hz AM noise (McFadden et al., 2013, 2014). In general, when listening to 40-Hz AM noise, EEG signals emanating from a broad area of the brain are synchronized at around 40 Hz (McFadden et al., 2014; Rauschecker & Scott, 2009; Reyes et al., 2004; Ross et al., 2005). Our method could successfully detect changes in dynamical structures when changes in dynamical structures of the network occurred with regards to the ASSR. This indicates that a model-based approach to detect the change point of functional brain networks is more convenient to interpret the temporal dynamics of the brain. Moreover, another remarkable point is that our method can detect changes in the dynamical structures reflected in scalp EEGs on a single-trial basis.

In summary, our proposed method successfully detected the dynamical structures of the brain networks, as well as the change points of those dynamical structures. Considering the increasing interest of neuroscientists in the relationship between flexible changes in functional brain networks and cognitive functions (Hutchison et al., 2013; Bassett et al., 2011; Baum et al., 2017; Braun et al., 2015, 2016), our method could help to reveal the crucial role of flexible changes in dynamical structures of the brain.

### 4.3. Significance of the proposed method in the context of change point detection

The change point is defined according to the model parameter *K* regarding the probability density *p_K_*(*x*) of the time series *x* changes from *K* = *K*_0_ to *K* = *K*_1_ (i.e., *K*_0_ ≠ *K*_1_), and the task of detecting the timing of such changes is called the change point detection problem. Change point detection has been actively studied in machine learning and data mining research over the past few decades (Basseville et al., 1993). Most prior methods of change point detection have been developed according to the common framework that the extent of abrupt changes in the time series data is measured using the log-likelihood ratio to compare the past and the current state of the probability density (Basseville et al., 1993). For the typical approach based on the log-likelihood ratio, the following methods can be considered: the cumulative sum of log-likelihood ratio algorithm (Basseville et al., 1993; Gustafsson & Gustafsson, 2000), the generalized log-likelihood ratio (Basseville, 1988; Gustafsson, 1996; Gustafsson & Gustafsson, 2000), and the AR model-based approach (Basseville, 1988; Yamanishi & Takeuchi, 2002). However, these typical methods are specialized for a single sequence, and a method that focuses on the time-evolving dynamics of network connectivity is still lacking. In a recent study, Idé and colleagues (Idé et al., 2009; Idé, 2014) proposed an approach for detecting structural changes in the network sequence based on the sparse Gaussian graphical model (graphical lasso), but this method estimates the graph structure assuming that there is a linear correlation between each node in the network. Therefore, this method is difficult to apply for multivariate time series data in which there is an underlying nonlinear dynamical system, such as synchronous neural activity data. As mentioned in the previous discussion (see section 4.1), in the field of bioinformatics, Xiong & Zhou (2013) proposed a Kalman filter-based change point detection algorithm for the time-varying gene regulatory network. However, this algorithm also assumes a linear model for temporal dynamics in the network. Moreover, as stated in section 4.1, the AR model-based method led to several false detections when applying it to the change point detection problem for nonlinear time-series data of a dynamical network such as the brain. Comparing our proposed method with those of the above-mentioned prior studies, our method also has the limitation that it applies a parametric model to estimate the network structure; however, our method has the advantage in that it can simultaneously estimate both visualization of the network structure and detection of temporal changes in such a network. Applying sequential Bayesian inference to estimate the model parameter is a common approach to model fitting. However, our method is unique in measuring the extent of the changes in model parameters by comparing prior and posterior distributions of model parameters for quantification of the change point score.

### 4.4. Limitations of this study

Despite the advantages of our proposed method, some limitations exist. First, the expressive power of phase dynamics in our method is strongly dependent on the pre-defined function of the dynamical model. Since phase dynamics are approximated by the Fourier series in our method, prediction performance and reliability depend on the dimension of the Fourier series. Second, the detection accuracy is also affected by the precision factor *β*, which was fixed at *β*^-1^ = 0.0001 in this study. As can be seen in Eq. (7), the parameter *β* affects the observed noise in the model. Given that ***f* Σ_K_f^T^**, which is contained in the right-hand side of Eq. (7), tends to be around zero throughout the sequential update of the predictive distributions, the prediction error covariance of the model converges with *β*^-1^*I*. This tendency directly influences the variance of the fitting error of the model. Thus, both the dimension of the Fourier series and *β* affect the performance of change point detection. However, these parameters should be selected exploratorily, because the optimal setting for each parameter depends on the complexity and dimensionality of the data. The optimal dimension of the Fourier series *P* could be estimated using the Akaike information criterion or Bayesian information criterion. For simplification, in this study, even though the dimension of the Fourier series is fixed at *P* = 1, future work should consider the dimensionality using the above-mentioned information criteria when applying the extended version of the proposed method. Moreover, to more strictly consider the effects of observation noise, applying a variational Bayesian linear regression approach with the Gamma prior distribution for the observation noise covariance (Tzikas et al., 2008) would enable us to deal with this issue. However, further studies are required to attempt this approach.

An additional limitation of our method is the number of oscillators required to guarantee an accurate estimation of the network couplings. As we described in the Supplementary Materials, the practical number of oscillators that could be accurately estimated by the proposed method is less than or equal to 10. We speculate that the reasons for this limitation are related to the limited dimension of the Fourier series *P* and noise estimation in Bayesian inference. Thus, the above-mentioned consideration regarding the improvements of these issues would be also helpful to increase the practical number of oscillators that our method could be applied to. The conventional methods to estimate the time-varying functional connectivity, such as correlation-based and VAR model-based methods, have no such limitation; however, these methods could lead to the incorrect detection of network couplings, as mentioned above.

Despite these limitations, our method provides advantages over the conventional methods in that both visualization of the network structure and detection of the change point are realized in parallel using Bayesian inference.

## 5. Conclusions

In this study, we tested a new method for detecting changes in dynamical structures of the brain. By applying our proposed method to numerical and electroencephalographic data, we confirmed that our method can successfully detect the change points at which dynamical changes of the network occur. These results suggest that our proposed method could help to reveal the neural basis of dynamical changes of brain networks. Furthermore, although it was beyond the scope of the current study, our proposed method could be applied to examine cognitive functions and learning capacity in more detail, as these functions are reflected in the time-evolving dynamics of brain networks. To apply our proposed method in future neuroscience studies, future studies are required to address some of the limitations mentioned above.

## 6. Acknowledgements

This research was funded by the JSPS KAKENHI grant (20K19867, https://kaken.nii.ac.jp/en/grant/KAKENHI-PROJECT-20K19867/) from the Japan Society for the Promotion of Science.

## 7. Credit author statement

Conceptualization, H.Y. and K.K.; Methodology, H.Y.; Software, H.Y.; Validation, H.Y. and K.K.; Formal analysis, H.Y.; Investigation, H.Y.; Resources, H.Y. and K.K.; Data curation, H.Y.; Writing-original draft preparation, H.Y.; Writing–review and editing, H.Y. and K.K.; Visualization, H.Y.; Supervision, K.K.; Project administration, K.K.; Funding acquisition, H.Y. All authors have read and agreed to the published version of the manuscript.

## 8. Data and code availability statements

Data will be made available to all interested researchers upon request. The sample code of the numerical simulation described in this paper is available at Github: https://github.com/myGit-YokoyamaHiroshi/ChangeDetectSim

## Supplementary Materials

### S1. Detailed description of mathematics in our proposed method

We will provide the theoretical background for the phase-coupled oscillator model and the details of our proposed method.

#### S1.1. Dynamical systems theory of the phase oscillator model

From the point of view of dynamical systems, the time series 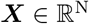, which is followed by a periodic function *f*, can be described as follows:

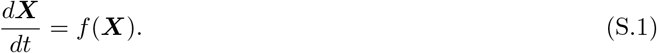

If the system exhibits limit cycles with stable point **X** = **X**_*τ*_(*ϕ*), the polar coordinates of this system can be described as follows:

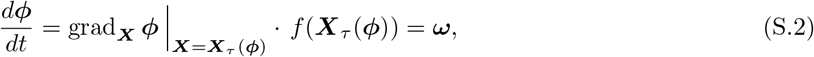

where *ϕ*, *ω*, and *τ* denote the phase, natural frequency, and one-cycle period, respectively. Phase dynamics follow the periodic cycle, which is described as *ϕ*(*t* + *τ*) = *ϕ*(*t*), around the stable point.

Here, we consider two coupled oscillators, ***X*** = {*x_i_*, *x_j_*}, driven by an interaction between oscillators. In this case, the dynamics can be described as follows:

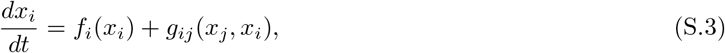

where *f_j_*(*x_j_*) and *g_j_*(*x_j_*, *x_i_*) indicate the dynamical system function of the *i*-th oscillator and the interaction between the *i* and *j*-th oscillators, respectively. In the polar coordinates, Eq. (S.3) can be expanded around the stable point *X_τ_*(Φ) = {*x_τ_*(*ϕ_i_*), *x_τ_*(*ϕ_j_*)}, as described in the following equation:

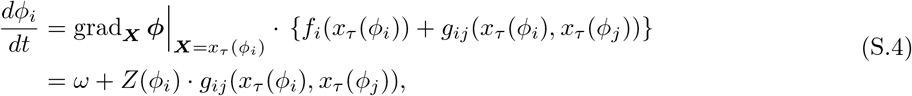

where 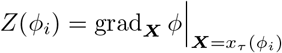 indicates the function of the phase-response curve corresponding to the *i*-th oscillator.

By introducing relative phase *ψ_i_* via *ϕ_i_* = *ωt* + *ψ_i_*, this can be rewritten as follows:

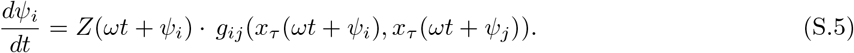

Moreover, by averaging over one cycle with the period *T*, this can be rewritten as follows:

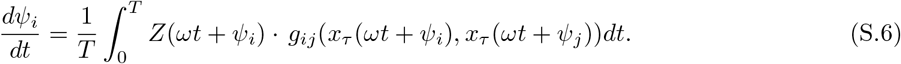

If *ωt* = *θ* is substituted to obtain 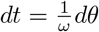, then 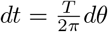. Therefore, we obtain the following equation:

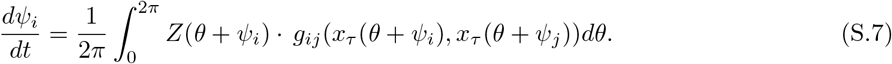

In addition, assuming that Θ = *θ* + *ψ_i_* (i.e., *θ* = Θ – *ψ_i_*), Eq. (S.5) can be approximated as the function of phase difference *ψ_j_* – *ψ_i_* (Kuramoto, 1984; Pietras & Daffertshofer, 2019), described as follows:

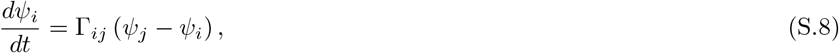

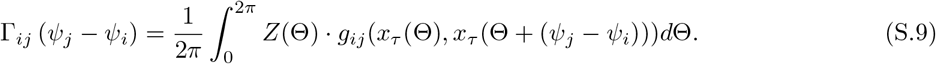

Note that the function Γ_*ij*_(*ψ_j_* – *ψ_i_*) describes the interaction between the *i* and *j*-th oscillators.

Considering Equations (S.5)–(S.9), Eq. (S.4) can be extended as the function of phase difference between oscillators (Kuramoto, 1984; Pietras & Daffertshofer, 2019), as follows:

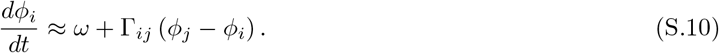

Given the above equations, the function Γ_*ij*_(*ϕ_j_* – *ϕ_i_*) describes the phase changes in the cycle period of coupled oscillators via network interactions for each oscillator. Thus, by estimating the function Γ_*ij*_(*ϕ_j_* – *ϕ_i_*) from empirically measured data, we can characterize the interaction between oscillators. In turn, by estimating the function Γ_*ij*_(*ϕ_j_* – *ϕ_i_*) from empirically measured data, we can characterize the interaction between oscillators. Hereafter, we call this function Γ_*ij*_(*ϕ_j_* – *ϕ_i_*) the “phase interaction function” (Penny et al., 2009; Ota & Aoyagi, 2014; Suzuki et al., 2018; Onojima et al., 2018).

#### S1.2. Phase oscillator model approximated with the Fourier series

Considering the phase oscillator model theory described in section S1.1, we applied the *N*-th phase-coupled oscillator model using the following equation to describe the dynamical network model in the brain:

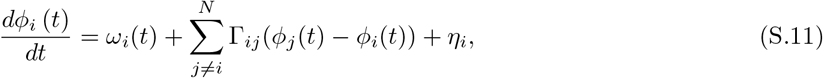

where *ϕ_i_*(*t*), *ω_i_*(*t*), and *η_i_*(*t*) indicate the phase, natural frequency, and observation noise of the i-th oscillator, respectively. In our study, as mentioned in the main manuscript, Eq. (S.11) was approximated using the *P*-th Fourier series as follows:

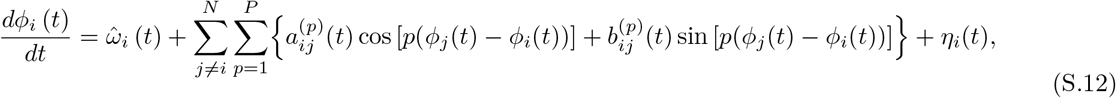

where

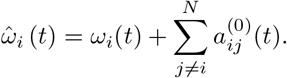

Note that 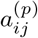 and 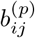 indicate the coefficients of the *p*-th Fourier series corresponding to the coupling strength between the *i* and *j*-th oscillators. Moreover, 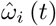 is the neutral frequency of this model. In our proposed method, the coupling strength *C_ij_* for all *i*, *j* is defined as follows:

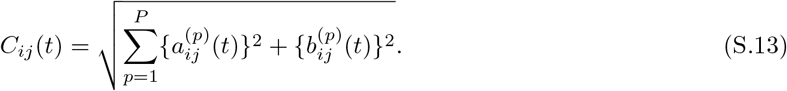

#### S1.3. Estimation of the model parameters and change point score

We will next explain how to predict the model parameters shown in Eq. (S.12) and the main idea behind change point detection in the observed data. To evaluate the temporal changes in the networks, we extended the parameter estimation algorithm in Eq. (S.12) using Bayesian inference (Bishop, 2007; Sarris, 1973). In this way, the parameters of Eq. (S.12) could be directly estimated from the observed data in the same way as for sequential Bayesian regression. To apply Bayesian inference, Eq. (S.12) was rewritten in its matrix form, as shown in Eq. (S.14).

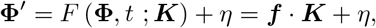

where,

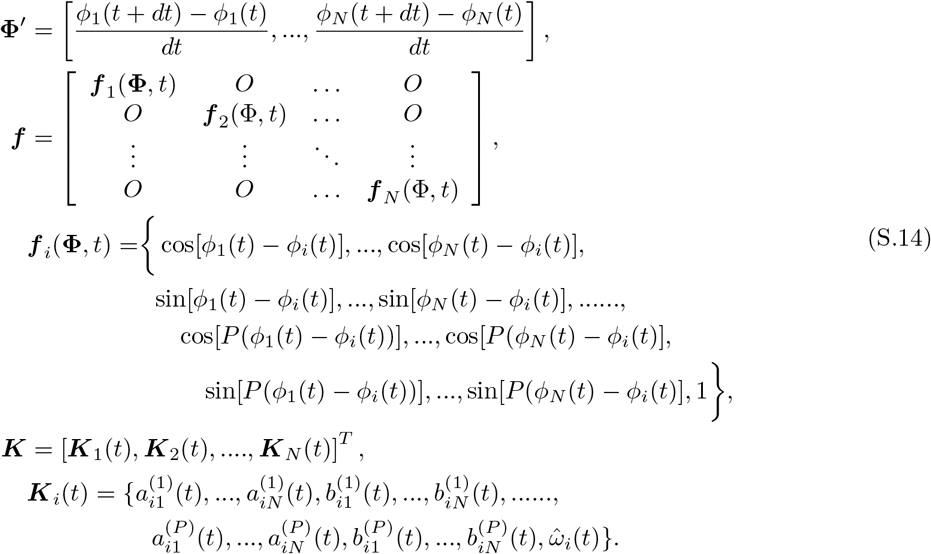

Note that *F* (**Φ**, *t*; ***K***) is the function of phase-coupled network dynamics in time *t*, characterized by phase vector *ϕ* and the parameter vector ***K***. In our model, *F* (**Φ**, *t*; ***K***) and ***K*** indicate the Fourier series and its coefficient, respectively. Moreover, **Φ′** indicates a phase velocity with a smaller time step *dt*. In the numerical simulations, we set the time step as *dt* = 0.01. For the real measured data, the time step *dt* corresponds to the sampling interval of the data.

Given the prior distribution of the model parameters, the parameter ***K*** in Eq. (S.14) can be calculated using the Bayesian rule (Bishop, 2007; Sarris, 1973), as follows:

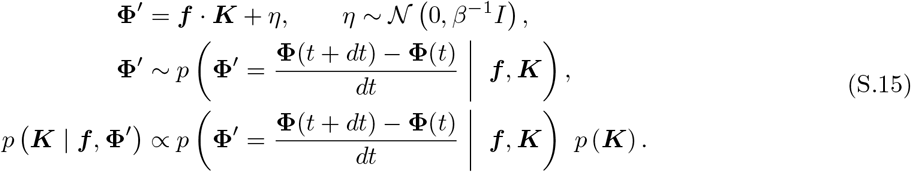

Here, in our proposed method, the prior distribution *p*(***K***) is assumed to have a multivariate normal distri-bution 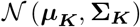, where ***μ_K_*** and **Σ_K_** indicate the mean and covariance of the distribution, respectively. The parameter *β* is a precision factor of the observation noise, and this parameter is an arbitrary constant term. In this study, this parameter was fixed as *β*^-1^ = 0.0001 for all analyses.

The update rule of the prior distribution is derived as follows (Sarris, 1973):

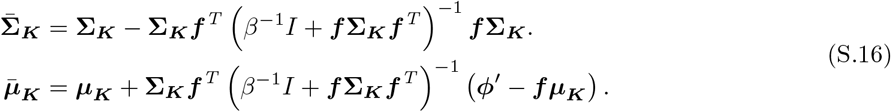

where 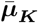 and 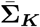 indicate the updated mean and covariance, respectively. After the update of the prior distribution, these parameters are set as 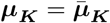 and 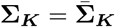 for the next updating process.

The posterior predictive distribution of the model can be inferred using the following equation:

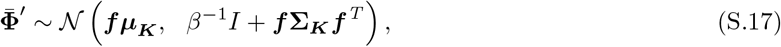

where 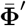 indicates the predicted phase velocity using the inferred model parameters. Based on the above procedure, the temporal changes in networks were estimated based on sequential Bayesian inference.

As a side note, by introducing the matrix ***G*** = ***Σ_K_f^T^***(*β*^-1^*I* + ***fΣ_K_f^T^***)^-1^ = ***Σ_K_f^T^S***^-1^ and the covariance ***S*** = *β*^-1^*I* + ***f Σ_K_ f^T^***, Eq. (S.16) can be rewritten as follows:

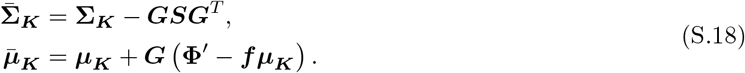

This matrix ***G*** is called the Kalman gain. Therefore, the sequential form of Bayesian regression with the multivariate normal prior is a special case of the Kalman filter (Sarris, 1973).

The advantage of applying Bayesian inference to estimate the function Γ_*ij*_(*ϕ_j_* – *ϕ_i_*) of dynamical networks is that the current state of the network dynamics and its dynamical structure can be expressed using a probability distribution. Therefore, our proposed method used the KL divergence between the prior and posterior distribution as an index of the change in dynamical structures, as shown in Eq. (S.19), and the value of this KL divergence is defined as the change point score (CPS), as follows:

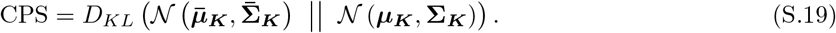

where ***μ_K_*** and 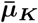 indicate the mean of the parameter *K* for the prior and posterior distribution, respectively. Also, **Σ_*K*_** and 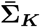 indicate the covariance of the parameter *K* for the prior and posterior distribution, respectively. Using the above process, we can detect change points in network dynamics while sequentially estimating dynamical structures in the brain using the KL divergence value. The summary of these procedures in our proposed method is shown in Fig. S1.

**Figure S1:**
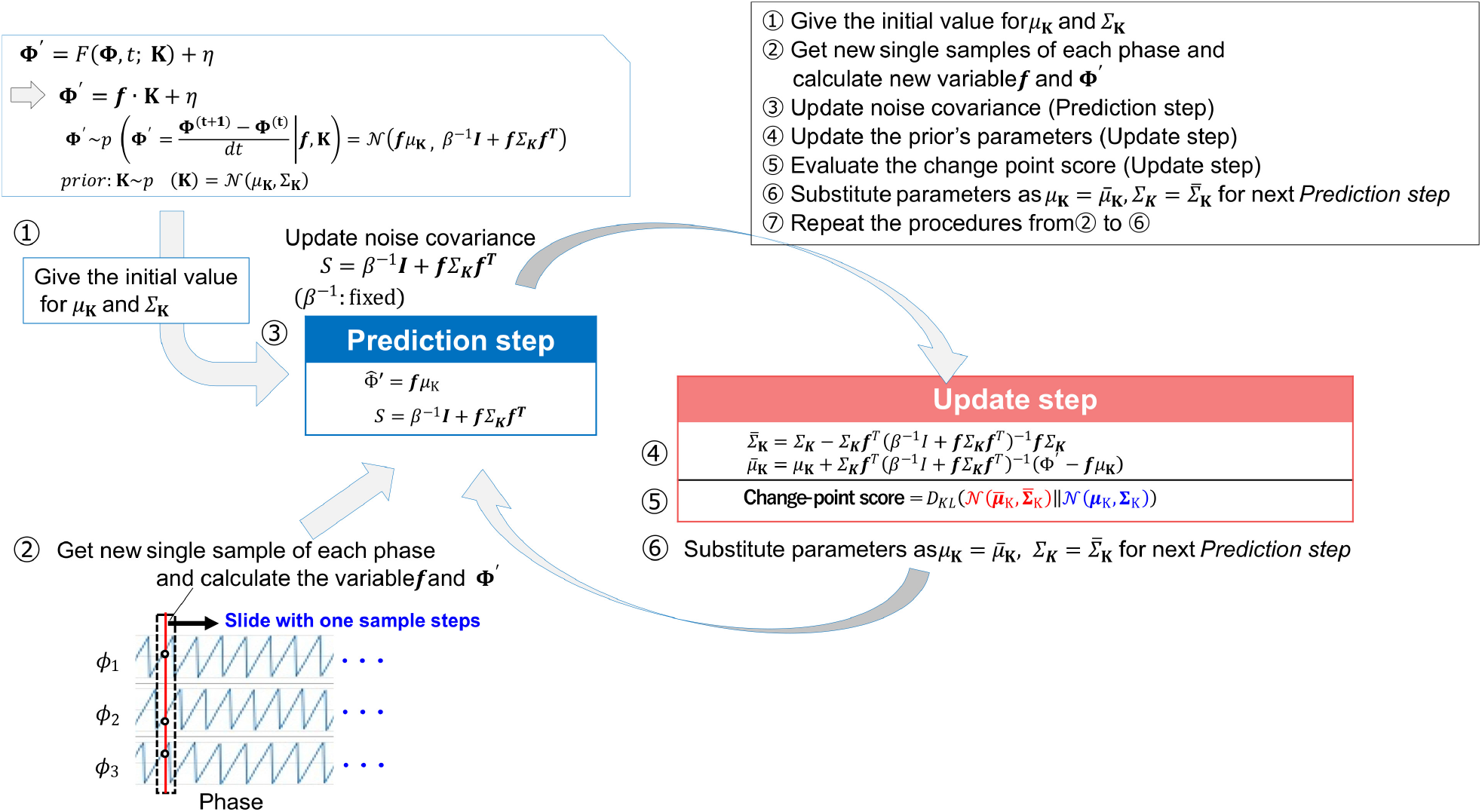
Schematic view of the sequential Bayesian inference algorithm. First, after giving the initial values of *μ_K_* and ∑_*k*_ for the prior distribution, the predictive posterior distribution of the model is updated with the sequentially sampled data *f* and **Φ′**(“Prediction step”), and the noise covariance *S* is updated. Next, the prior distribution of the model parameters is updated (“Update step”). After this, the extent of change in dynamical structures of the network is assessed using the Kullback-Leibler divergence (i.e., evaluation of the change point score). The updated parameters of the prior distribution are set as 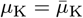 and 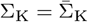 for the next “Prediction step”. According to the above-mentioned description, the model parameter updating process is conducted on a sample-by-sample basis, and the metric of change point score using the KL divergence is also calculated after updating the prior of the model parameters sample-by-sample.

#### S1.4. Theoretical background of change detection problems

As mentioned in the section S1.3, in our proposed method, the extent of change in the network structures is quantified based on the comparison between prior and posterior distributions of model parameters. From the viewpoint of change point detection based on probability statistics, if the past and current state of variable *x* can explain the probability density *p*(*x*) and *q*(*x*), the changes in variable *x* are typically measured as the gap between *p*(*x*) and *q*(*x*) using the log-likelihood ratio (Basseville et al., 1993). Moreover, the expectation under the probability *p*(*x*) of the log-likelihood ratio between *p*(*x*) and *q*(*x*) can be described as the KL divergence (Basseville et al., 1993), as follows:

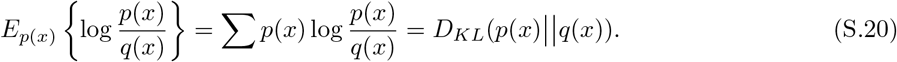

Note that *E*_*p*(*x*)_{·} indicates the expectation operator under the probability *p*(*x*). *D_KL_*(*p*(*x*)ļļ*q*(*x*)) indicates the KL divergence from *q*(*x*) to *p*(*x*). Based on these statistical backgrounds, we measured the extent of change in the network structures using the KL divergence between the prior and posterior probability density of the model parameters as in Eq. (S.19).

### S2. Generation of synthetic data

In all simulations, which are described in the main manuscript and supplementary sections S3.1 and S3.2, we used the stochastic phase-coupled oscillator model approximated with a first-order Fourier series to generate the synthetic data. The model used is defined as follows:

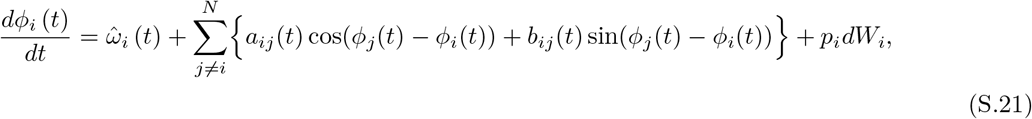

where

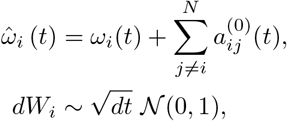

where *p_i_* and *dW_i_* indicate the noise scaling factor and white noise factor, which followed a normal distribution 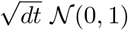. The synthetic data were generated by the numerical integration of Eq. (S.21) using the Euler-Maruyama method. The integration scheme of the Euler-Maruyama method in Eq. (S.21) is repetitively conducted using the following equation:

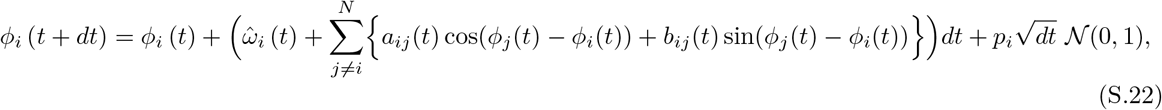

where *dt* indicates the time step. As mentioned above, in the numerical simulations, the time step was set as *dt* = 0.01. Other parameters, such as the Fourier coefficient *a_ij_*, *b_ij_*, natural frequency 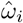, and noise-scaling factor *p_i_*, were set as the different values depending on the settings for each simulation.

### S3. Supplementary simulations

As stated in the main manuscript, we conducted three different numerical simulations; only one of these simulations is reported in the main manuscript. Here, we will provide the detailed descriptions and results for the remaining two simulations.

#### S3.1 Supplementary simulation A

##### S3.1.1. Simulation settings

The purpose of this simulation was to clarify the number of practical oscillators and the number of learning iterations of the model required to accurately estimate the network couplings when using our proposed method. To validate this, we applied our proposed method to synthetic time-series data generated by *N_oscj_*-coupled oscillators with fixed network couplings.

In this simulation, we set four conditions of *N_oscj_* = [3, 10, 15, 20, 30], and generated 4,000 samples of synthetic data for each condition of *N_oscj_* using Eq. (S.22). The parameter settings of Fourier coefficients *a_ij_* and *b_ij_* for each condition were selected, as shown in Figure S2. Moreover, the natural frequency 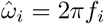 of the *i*-th oscillator is set as a fixed value so that *f_i_* follows a normal distribution 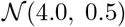. The noise-scaling factor *p_i_* for the integration scheme in Eq. (S.22) was set as *p_i_* = 0.001.

After generating the synthetic data using Eq. (S.22) with the above parameter settings, our Bayesian inference-based method was applied to the time series of synthetic phase data. The precision parameter *β* in Eqs. (S.16) and (S.17) was fixed at *β*^−1^ = 0.0001 for this simulation. For each iteration of the model updating steps, we evaluated the error of the structure of network couplings with the mean absolute error (MAE) to evaluate the estimation accuracy, as follows:

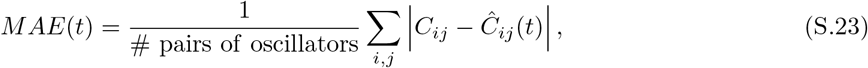

where *C_ij_* indicates the exact coupling strength value between the pair of oscillators *i, j* in the synthetic data, and *Ĉ_ij_*(*t*) indicates the estimated value of coupling strength between the *i*-th and *j*-th oscillators in time *t*. The value of *Ĉ_ij_*(*t*) in time *t* was calculated using estimated Fourier coefficients *â_ij_*(*t*) and 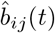 predicted by sequential Bayesian inference, as shown in Figure S1.

To calculate the confidence intervals of estimation error of the coupling strength *Ĉ_ij_*(*t*), the surrogate method was applied. Specifically, to generate the null distribution of the randomized MAE, we first calculated surrogate Cj(t), in which the temporal order of the estimated *Ĉ_ij_*(*t*) shuffled 1,000 times, and the MAE between the true network coupling strength *C_ij_* and the shuffled *Ĉ_ij_*(*t*) was calculated. Then, the resulting values of these surrogate MAEs were used to compose the null distribution of MAEs, and to assess the 95% confidence intervals of the estimation error.

**Figure S2:**
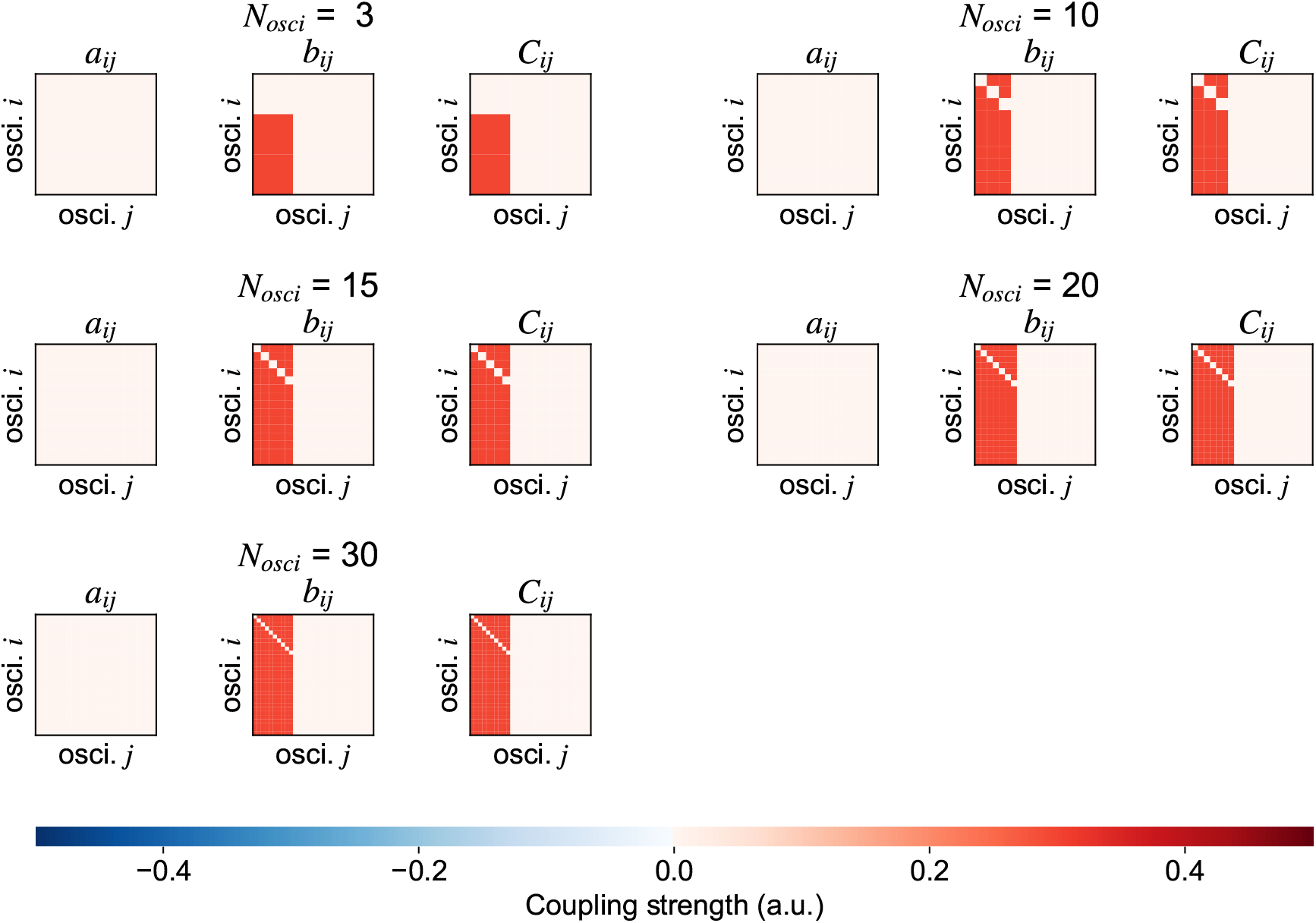
Parameter settings in numerical simulation A. This simulation set five different conditions for the number of oscillators (*N_Osci_* = 3, 10, 15, 20, and 30). For each condition, the time series of synthetic data in this simulation were generated by a first-order Fourier series-approximated phase-coupled oscillator model with the parameter settings shown in this figure. The Fourier coefficients *a_i,j_* and coupling strength *C_ij_* were selected such that one-third of oscillator pairs interacted.

##### S3.1.2. Results

The results of this simulation are shown in Figure S3A, which shows the changes in the estimated network couplings for each learning iteration. For less than *N_osci_* = 10, the estimated structures of the network couplings became close to the exact structure of the synthetic data after 1,000 parameter updating iterations (see iteration 1,000 in Figure S3A). However, *N_osci_* ≥ 15 required over 2,000 iterations to accurately estimate the network couplings.

To confirm these results in more detail, we evaluated changes in the estimated error for each learning iteration with all oscillator number conditions (Figure S3B). As a result, the errors decreased to around zero in conditions with less than *N_osci_* = 10 at 1,000 iterations. By contrast, for *N_osci_* ≥ 15, although the error finally decreased to around zero, the iteration number required to accurately estimate the network couplings gradually increased in proportion to the increase in the number of oscillators.

Furthermore, to confirm the number of learning iterations required to accurately estimate the network couplings, the error for each condition was compared with the 95% confidence intervals, which were evaluated using surrogate error data for each condition. The results revealed that, when applied to time series data with less than *N_osci_* = 10, our proposed method can ensure the accurate estimation of network couplings with less than 500 model parameter updating iterations.

Considering these results for this simulation, the practical number of oscillators that could be applied our method to guarantee estimation accuracy is less than *N_osci_* = 10. Therefore, for the following analyses, we applied our method to the time series data within the number of oscillators (or dimension of time series data) N_*osci*_ = 10.

**Figure S3:**
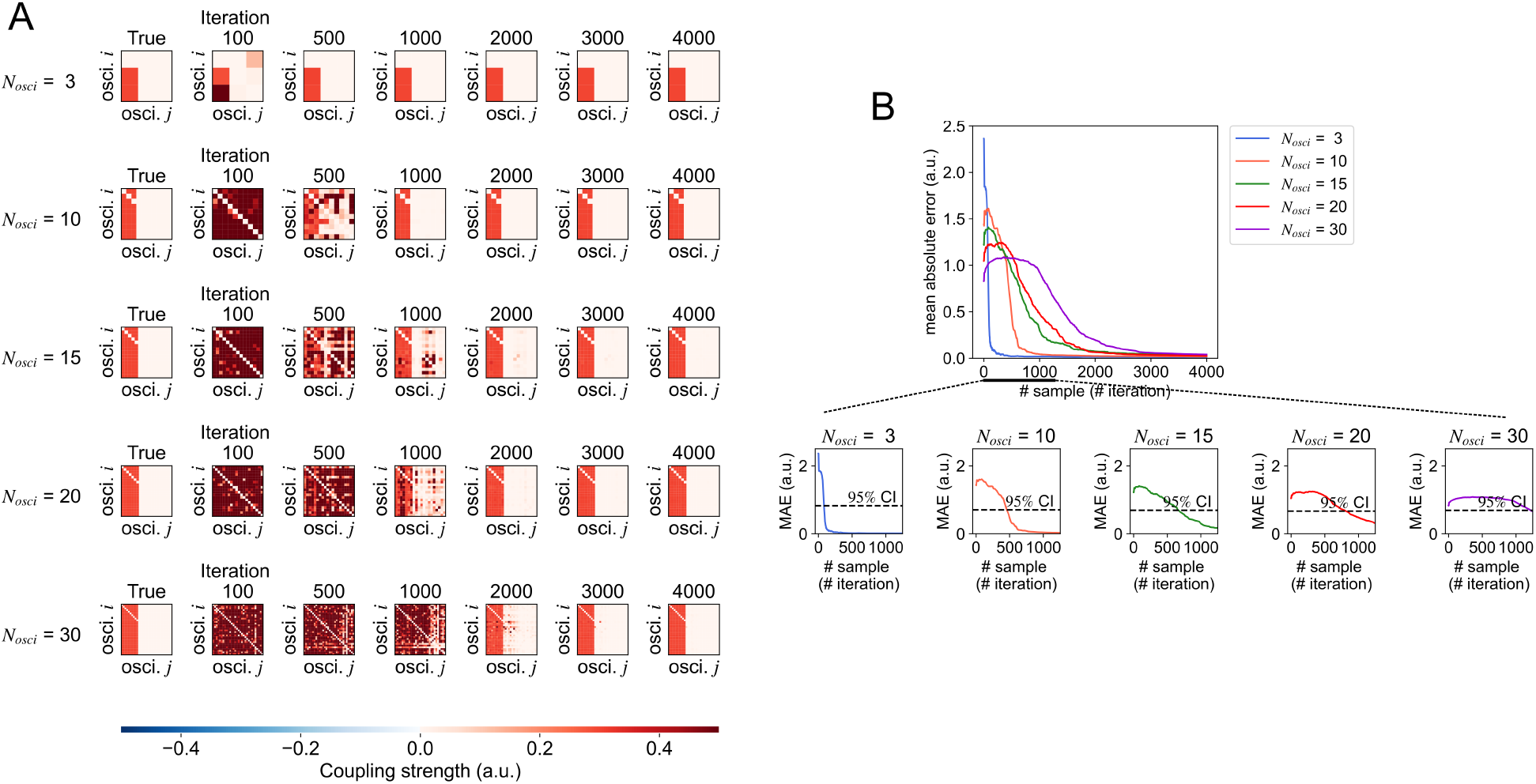
Comparison of true and predicted network structures in numerical simulation A. (A) The changes in the estimated network strength *Ĉ_ij_* for each iteration (note: an iteration indicates the number of updates in the model with sequential Bayesian inference). (B) Comparison of changes in the error of network structures for each oscillator number condition.

#### S3.2. Supplementary simulation B

##### S3.2.1. Simulation settings

To clarify whether our method could sequentially detect the changes in network couplings, we applied our method to 3,000 samples of synthetic time series data of ten coupled oscillators generated by Eq. (S.22) with the first-order Fourier series. When generating the synthetic data, the noise-scaling factor *p_i_* in Eq. (S.22) was fixed as *p_i_* = 0.001 for each *i*-th oscillator. These synthetic time series consisted of three segments, and the parameters *a_ij_* and *b_ij_* changed for each segment. The parameter setting for *a_ij_* and *b_ij_* is shown in Figure S4.

**Figure S4:**
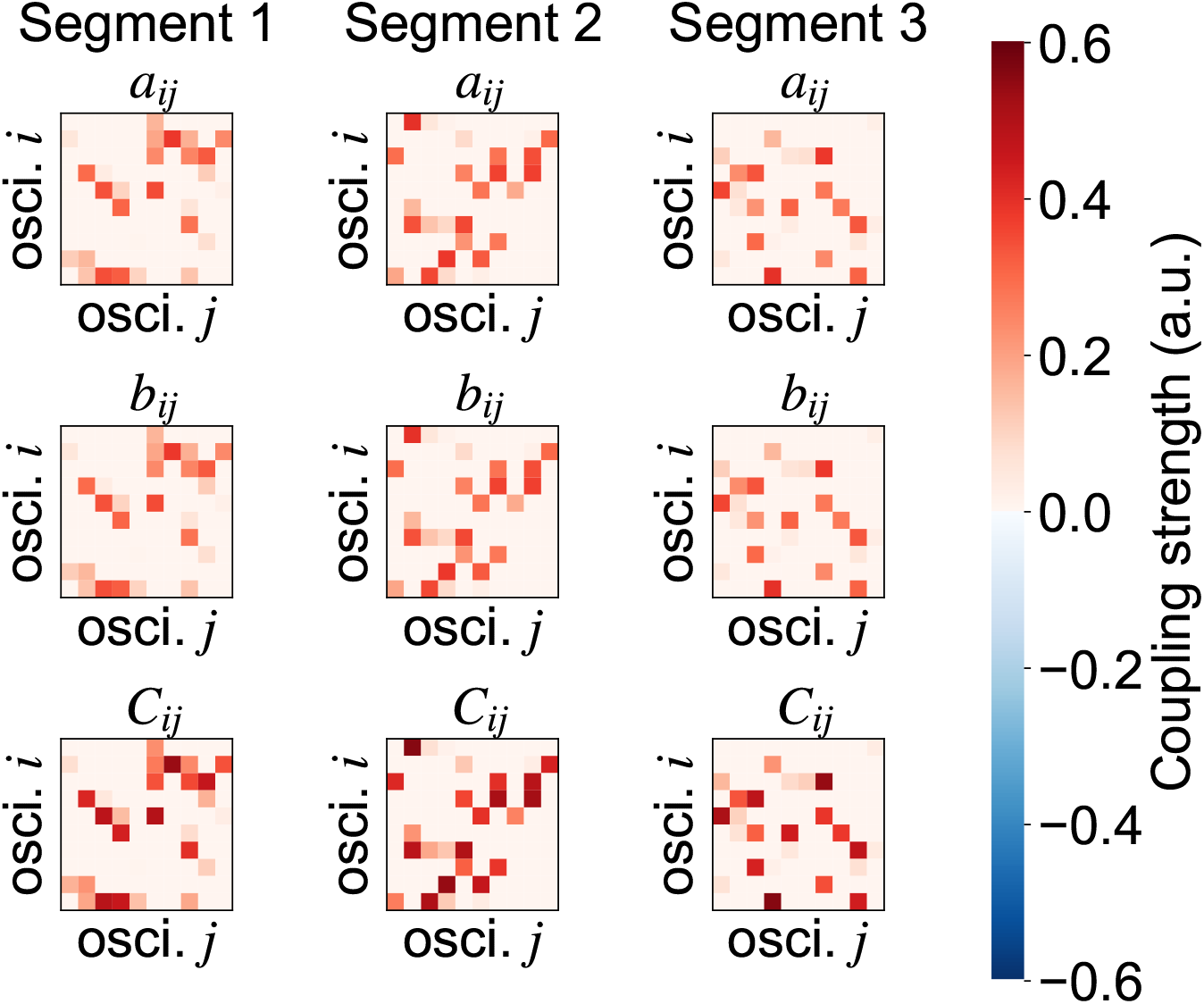
Parameter settings in numerical simulation C. The synthetic data of ten oscillators were generated by the first-order Fourier series-approximated phase-coupled oscillator model with these parameter settings. The generated synthetic time series consisted of three segments separated by two events.

After generating the synthetic data using the above-mentioned procedures with the parameter settings shown in Figure S4, our method was applied to these synthetic data as follows:

- The time series of synthetic data was fit to the first-order Fourier series using Bayesian inference
- The KL divergence between prior and posterior model parameters was evaluated (i.e., we calculated the change point score)
- The time-sample with a greater change point score relative to the threshold *ζ* was selected as the “change point”

In this simulation, the threshold *ζ* for the change point score was set as the mean(KL) + 3SD(KL); these terms refer to the mean KL divergence and 3 standard deviations of the KL divergence of the change point score across samples, respectively. The precision parameter *β* in Eqs. (S.16) and (S.17) was fixed at *β*^−1^ = 0.0001 for this simulation.

##### S3.2.2. Results

The result for this simulation is shown in Figure S5. As can be seen in Figure S5B, a significant increase of the change point score was detected around the boundary of each segment (see the red dot in Figure S5B). Moreover, as shown in Figure S5A, the predicted network couplings of each segment were relatively similar to the true network couplings in the synthetic data for each segment. These results indicated that our method could also be applied for ten coupled oscillators as the detector of the change point of dynamical structure in phase-coupled oscillator networks.

**Figure S5:**
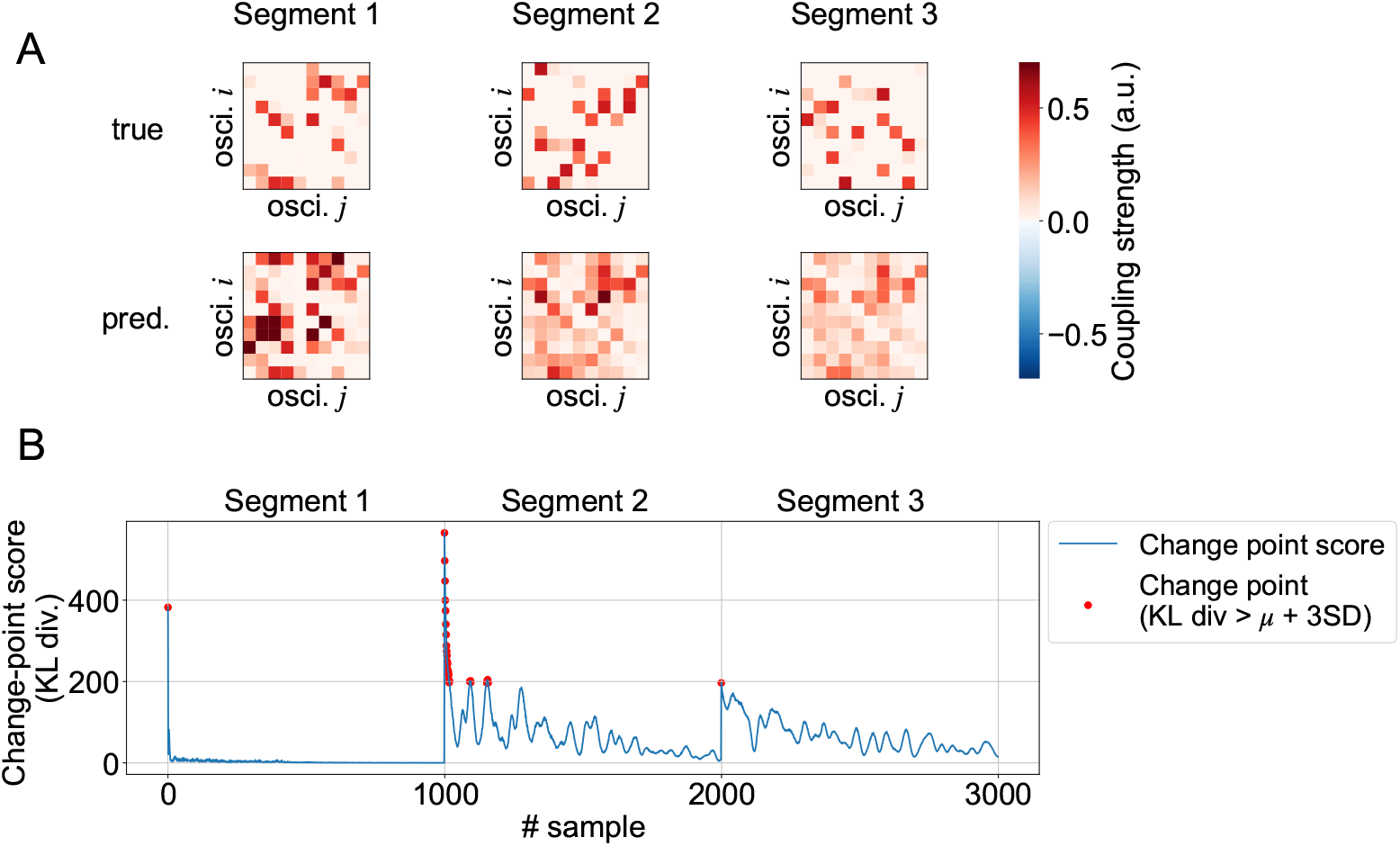
Change point detection in numerical simulation C. (A) Comparison of true and estimated network couplings. The upper three panels show the true network couplings in each segment. The lower three panels show the estimated network couplings in each segment estimated by our method. (B) The time course of the change point scores. The red dots indicate the selected change points based on the pre-defined threshold.

## Data and code availability statements

The sample code of the numerical simulation described in this supplementary material is available at Github: https://github.com/myGit-YokoyamaHiroshi/ChangeDetectSim/tree/master/supplementary

## Notes

### Competing Interest Statement

The authors have declared no competing interest.

## References

Astolfi, L., Cincotti, F., Mattia, D., De Vico Fallani, F., Tocci, A., Colosimo, A., Salinari, S., Marciani, M., Hesse, W., Witte, H., Ursino, M., Zavaglia, M., & Babiloni, F. (2008). Tracking the Time-Varying Cortical Connectivity Patterns by Adaptive Multivariate Estimators. IEEE Transactions on Biomedical Engineering, 55, 902–913. URL: http://ieeexplore.ieee.org/document/4360119/. doi:10.1109/TBME.2007.905419.

Bassett, D. S., Wymbs, N. F., Porter, M. A., Mucha, P. J., Carlson, J. M., & Grafton, S. T. (2011). Dynamic reconfiguration of human brain networks during learning. Proceedings of the National Academy of Sciences of the United States of America, 108, 7641–7646. doi:10.1073/pnas.1018985108.

Basseville, M. (1988). Detecting changes in signals and systems—A survey. Automatica, 24, 309–326. URL: https://linkinghub.elsevier.com/retrieve/pii/0005109888900738. doi:10.1016/0005-1098(88)90073-8.

Basseville, M., Nikiforov, I. V. et al. (1993). Detection of abrupt changes: theory and application volume 104. Prentice hall Englewood Cliffs.

Baum, G. L., Ciric, R., Roalf, D. R., Betzel, R. F., Moore, T. M., Shinohara, R. T., Kahn, A. E., Vandekar, S. N., Rupert, P. E., Quarmley, M., Cook, P. A., Elliott, M. A., Ruparel, K., Gur, R. E., Gur, R. C., Bassett, D. S., & Satterthwaite, T. D. (2017). Modular Segregation of structural brain networks supports the development of executive function in youth. Current Biology, 27, 1561–1572.e8. URL: http://dx.doi.org/10.1016/j.cub.2017.04.051. doi:10.1016/j.cub.2017.04.051.

Bishop, C. M. (2007). Pattern recognition and machine learning (Information science and statistics). (1st ed.). Springer.

Braun, U., Schäfer, A., Bassett, D. S., Rausch, F., Schweiger, J. I., Bilek, E., Erk, S., Romanczuk-Seiferth, N., Grimm, O., Geiger, L. S., Haddad, L., Otto, K., Mohnke, S., Heinz, A., Zink, M., Walter, H., Schwarz, E., Meyer-Lindenberg, A., & Tost, H. (2016). Dynamic brain network reconfiguration as a potential schizophrenia genetic risk mechanism modulated by NMDA receptor function. Proceedings of the National Academy of Sciences of the United States of America, 113, 12568–12573. doi:10.1073/pnas.1608819113.

Braun, U., Schäfer, A., Walter, H., Erk, S., Romanczuk-Seiferth, N., Haddad, L., Schweiger, J. I., Grimm, O., Heinz, A., Tost, H., Meyer-Lindenberg, A., & Bassett, D. S. (2015). Dynamic reconfiguration of frontal brain networks during executive cognition in humans. Proceedings of the National Academy of Sciences of the United States of America, 112, 11678–11683. doi:10.1073/pnas.1422487112.

Breakspear, M., Heitmann, S., & Daffertshofer, A. (2010). Generative models of cortical oscillations: neurobiological implications of the kuramoto model. Frontiers in Human Neuroscience, 4, 1–14. URL: http://journal.frontiersin.org/article/10.3389/fnhum.2010.00190/abstract. doi:10.3389/fnhum.2010.00190.

Cabral, J., Hugues, E., Sporns, O., & Deco, G. (2011). Role of local network oscillations in resting-state functional connectivity. NeuroImage, 57, 130–139. URL: http://dx.doi.org/10.1016/j.neuroimage.2011.04.010. doi:10.1016/j.neuroimage.2011.04.010.

Damaraju, E., Allen, E. A., Belger, A., Ford, J. M., McEwen, S., Mathalon, D. H., Mueller, B. A., Pearlson, G. D., Potkin, S. G., Preda, A., Turner, J. A., Vaidya, J. G., Van Erp, T. G., & Calhoun, V. D. (2014). Dynamic functional connectivity analysis reveals transient states of dysconnectivity in schizophrenia. NeuroImage: Clinical, 5, 298–308. URL: http://dx.doi.org/10.1016/j.nicl.2014.07.003. doi:10.1016/j.nicl.2014.07.003.

Deco, G., Cabral, J., Woolrich, M. W., Stevner, A. B., van Hartevelt, T. J., & Kringelbach, M. L. (2017a). Single or multiple frequency generators in on-going brain activity: a mechanistic whole-brain model of empirical MEG data. NeuroImage, 152, 538–550. doi:10.1016/j.neuroimage.2017.03.023.

Deco, G., Kringelbach, M. L., Jirsa, V. K., & Ritter, P. (2017b). The dynamics of resting fluctuations in the brain: metastability and its dynamical cortical core. Scientific Reports, 7, 1–14. doi:10.1038/s41598-017-03073-5.

Flandrin, P., Goncalves, P., & Rilling, G. (2004). Detrending and denoising with empirical mode decompositions. In European Signal Processing Conference. volume 06-10-Sept.

Flandrin, P., Goncalves, P., & Rilling, G. (2005). EMD equivalent filter banks, from interpretation to applications. (pp. 57–74). WORLD SCIENTIFIC volume 5 of Interdisciplinary Mathematical Sciences. URL: https://www.worldscientific.com/worldscibooks/10.1142/5862 http://www.worldscientific.com/doi/abs/10.1142/9789812703347{_}0003. doi:10.1142/9789812703347_0003.

Gensler, A., & Sick, B. (2014). Novel criteria to measure performance of time series segmentation techniques. CEUR Workshop Proceedings, 1226, 193–204.

Granger, C. W., Newbold, P., & Econom, J. (1974). Spurious regressions in econometrics. Baltagi, Badi H. A Companion of Theoretical Econometrics, 2, 557–61. URL: https://doi.org/10.1016/0304-4076(74)90034-7. doi:10.1016/0304-4076(74)90034-7.

Gustafsson, F. (1996). The marginalized likelihood ratio test for detecting abrupt changes. IEEE Transactions on Automatic Control, 41, 66–78. URL: http://ieeexplore.ieee.org/document/481608/. doi:10.1109/9.481608.

Gustafsson, F., & Gustafsson, F. (2000). Adaptive filtering and change detection volume 1. Citeseer.

Hutchison, R. M., Womelsdorf, T., Allen, E. A., Bandettini, P. A., Calhoun, V. D., Corbetta, M., Della Penna, S., Duyn, J. H., Glover, G. H., Gonzalez-Castillo, J., Handwerker, D. A., Keilholz, S., Kiviniemi, V., Leopold, D. A., de Pasquale, F., Sporns, O., Walter, M., & Chang, C. (2013). Dynamic functional connectivity: promise, issues, and interpretations. NeuroImage, 80, 360–378. URL: http://dx.doi.org/10.1016/j.neuroimage.2013.05.079. doi:10.1016/j.neuroimage.2013.05.079. arXiv:NIHMS150003.

Idé, T. (2014). Change Detection from Heterogeneous Data Sources. In Data Mining for Service (pp. 221–243). volume 3. URL: http://link.springer.com/10.1007/978-3-642-45252-9 http://link.springer.com/10.1007/978-3-642-45252-9_13. doi:10.1007/978-3-642-45252-9_13.

Idée, T., Lozano, A. C., Abe, N., & Liu, Y. (2009). Proximity-based anomaly detection using sparse structure learning. Society for Industrial and Applied Mathematics - 9th SIAM International Conference on Data Mining 2009, Proceedings in Applied Mathematics, 1, 96–107. doi:10.1137/1.9781611972795.9.

Kayser, J., Tenke, C. E., Gates, N. A., Kroppmann, C. J., Gil, R. B., & Bruder, G. E. (2006). Erp/csd indices of impaired verbal working memory subprocesses in schizophrenia. Psychophysiology, 43, 237–252.

Ko, T.-W., & Ermentrout, G. B. (2009). Phase-response curves of coupled oscillators. Physical Review E, 79, 016211. URL: https://link.aps.org/doi/10.1103/PhysRevE.79.016211. doi: 10.1103/PhysRevE.79.016211. arXiv:arXiv:0809.3371v1.

Kovécs, G., Sebestyen, G., & Hangan, A. (2020). Evaluation metrics for anomaly detection algorithms in time-series. Acta Universitatis S’apientiae, Informatica, 11, 113–130. doi:10.2478/ausi-2019-0008.

Kuramoto, Y. (1984). Chemical oscillations, waves, and turbulence. Springer. URL: hhttps://www.springer.com/gp/book/9783642696916. doi: 10.1007/978-3-642-69689-3.

Leistritz, L., Pester, B., Doering, A., Schiecke, K., Babiloni, F., Astolfi, L., & Witte, H. (2013). Time-variant partial directed coherence for analysing connectivity: A methodological study. Philosophical Transactions of the Royal Society A: Mathematical, Physical and Engineering Sciences, 371. doi:10.1098/rsta.2011.0616.

Lie, O. V., & van Mierlo, P. (2017). Seizure-Onset Mapping Based on Time-Variant Multivariate Functional Connectivity Analysis of High-Dimensional Intracranial EEG: A Kalman Filter Approach. Brain Topography, 30, 46–59. doi:10.1007/s10548-016-0527-x.

Liu, S., Yamada, M., Collier, N., & Sugiyama, M. (2013). Change-point detection in time-series data by relative density-ratio estimation. Neural Networks, 43, 72–83. doi:10.1016/j.neunet.2013.01.012.

McFadden, K., Steinmetz, S., Carroll, A., Simon, S., Wallace, A., & Rojas, D. (2013). EEG auditory steady state reliability paper,. URL: https://figshare.com/articles/dataset/EEG_Auditory_Steady_State_Reliability_Paper/829584. doi:10.6084/m9.figshare.829584.v14.

McFadden, K. L., Steinmetz, S. E., Carroll, A. M., Simon, S. T., Wallace, A., & Rojas, D. C. (2014). Test-retest reliability of the 40 Hz EEG auditory steady-state response. PLoS ONE, 9, 59–61. doi:10.1371/journal.pone.0085748.

Molla, M. K. I., Tanaka, T., Rutkowski, T. M., & Cichocki, A. (2010). Separation of EOG artifacts from EEG signals using bivariate EMD. In 2010 IEEE International Conference on Acoustics, Speech and Signal Processing (pp. 562–565). IEEE. URL: http://ieeexplore.ieee.org/document/5495594/. doi:10.1109/ICASSP.2010.5495594.

Netoff, T., Schwemmer, M. A., & Lewis, T. J. (2012). Phase response curves in neuroscience. URL: http://link.springer.com/10.1007/978-1-4614-0739-3. doi:10.1007/978-1-4614-0739-3.

Onojima, T., Goto, T., Mizuhara, H., & Aoyagi, T. (2018). A dynamical systems approach for estimating phase interactions between rhythms of different frequencies from experimental data. PLoS Computational Biology, (pp. 1–20). doi:10.1371/journal.pcbi.1005928.

Ota, K., & Aoyagi, T. (2014). Direct extraction of phase dynamics from fluctuating rhythmic data based on a bayesian approach. doi:10.1016/j.ceb.2004.02.009. arXiv:arXiv:1405.4126v1.

Penny, W. D., Litvak, V., Fuentemilla, L., Duzel, E., & Friston, K. (2009). Dynamic Causal Models for phase coupling. Journal of Neuroscience Methods, 183, 19–30. URL: http://dx.doi.org/10.1016/j.jneumeth.2009.06.029. doi:10.1016/j.jneumeth.2009.06.029.

Pietras, B., & Daffertshofer, A. (2019). Network dynamics of coupled oscillators and phase reduction techniques. Physics Reports, 819, 1–105. URL: https://doi.org/10.1016/j.physrep.2019.06.001. doi:10.1016/j.physrep.2019.06.001.

Rauschecker, J. P., & Scott, S. K. (2009). Maps and streams in the auditory cortex: nonhuman primates illuminate human speech processing. Nature Neuroscience, 12, 718–724. doi:10.1038/nn.2331.

Reyes, S. A., Salvi, R. J., Burkard, R. F., Coad, M. L., Wack, D. S., Galantowicz, P. J., & Lockwood, A. H. (2004). PET imaging of the 40 Hz auditory steady state response. Hearing Research, 194, 73–80. doi:10.1016/j.heares.2004.04.001.

Ross, B., Herdman, A. T., & Pantev, C. (2005). Right hemispheric laterality of human 40 Hz auditory steady-state responses. Cerebral Cortex, 15, 2029–2039. doi:10.1093/cercor/bhi078.

Sarris, A. H. (1973). A bayesian approach to estimation of time-varying regression coefficients. Journal of economic and social measurement, 2, 501–523. URL: http://www.nber.org/chapters/c9941.

Sase, T., & Kitajo, K. (2021). The metastable brain associated with autistic-like traits of typically developing individuals. PLOS’ Computational Biology, 17, e1008929. URL: http://dx.doi.org/10.1371/journal.pcbi.1008929. doi:10.1371/journal.pcbi.1008929.

Stankovski, T., Pereira, T., McClintock, P. V., & Stefanovska, A. (2017). Coupling functions: universal insights into dynamical interaction mechanisms. Reviews of Modern Physics, 89, 045001. URL: https://link.aps.org/doi/10.1103/RevModPhys.89.045001. doi:10.1103/RevModPhys.89.045001. arXiv:1706.01810.

Suzuki, K., Aoyagi, T., & Kitano, K. (2018). Bayesian estimation of phase dynamics based on partially sampled spikes generated by realistic model neurons. Frontiers in Computational Neuroscience, 11, 1–13. URL: http://journal.frontiersin.org/article/10.3389/fncom.2017.00116/full. doi:10.3389/fncom.2017.00116.

Tzikas, D. G., Likas, A. C., & Galatsanos, N. P. (2008). The variational approximation for Bayesian inference: Life after the EM algorithm. IEEE Signal Processing Magazine, 25, 131–146. doi:10.1109/MSP.2008.929620.

Xiong, J., & Zhou, T. (2013). A Kalman-Filter Based Approach to Identification of Time-Varying Gene Regulatory Networks. PLoS ONE, 8. doi:10.1371/journal.pone.0074571.

Yamanishi, K., & Takeuchi, J. I. (2002). A unifying framework for detecting outliers and change points from non-stationary time series data. Proceedings of the ACM SIGKDD International Conference on Knowledge Discovery and Data Mining, (pp. 676–681). doi:10.1145/775107.775148.

Ying, J., Zhou, D., Lin, K., & Gao, X. (2015). Network analysis of functional brain connectivity driven by gamma-band auditory steady-state response in auditory hallucinations. Journal of Medical and Biological Engineering, 35, 45–51. doi:10.1007/s40846-015-0004-0.

## Supplementary References

Kuramoto, Y. (1984). Chemical oscillations, waves, and turbulence. Springer. URL: hhttps://www.springer.com/gp/book/9783642696916. doi:10.1007/978-3-642-69689-3.

